# Mapping and modeling the genomic basis of differential RNA isoform expression at single-cell resolution with LR-Split-seq

**DOI:** 10.1101/2021.04.26.441522

**Authors:** Elisabeth Rebboah, Fairlie Reese, Katherine Williams, Gabriela Balderrama-Gutierrez, Cassandra McGill, Diane Trout, Isaryhia Rodriguez, Heidi Liang, Barbara J. Wold, Ali Mortazavi

**Author notes:** These authors contributed equally to this work (Elisabeth Rebboah, Fairlie Reese).

## Abstract

Alternative RNA isoforms are defined by promoter choice, alternative splicing, and polyA site selection. Although differential isoform expression is known to play a large regulatory role in eukaryotes, it has proved challenging to study with standard short-read RNA-seq because of the uncertainties it leaves about the full-length structure and precise termini of transcripts. The rise in throughput and quality of long-read sequencing now makes it possible, in principle, to unambiguously identify most transcript isoforms from beginning to end. However, its application to single-cell RNA-seq has been limited by throughput and expense. Here, we develop and characterize long-read Split-seq (LR-Split-seq), which uses a combinatorial barcoding-based method for sequencing single cells and nuclei with long reads. We show that LR-Split-seq can associate isoforms with cell types with relative economy and design flexibility. We characterize LR-Split-seq for whole cells and nuclei by using the well-studied mouse C2C12 system in which mononucleated myoblast cells differentiate and fuse into multinucleated myotubes. We show that the overall results are reproducible when comparing long- and short-read data from the same cell or nucleus. We find substantial evidence of differential isoform expression during differentiation including alternative transcription start site (TSS) usage. We integrate the resulting isoform expression dynamics with snATAC-seq chromatin accessibility to validate TSS-driven isoform choices. LR-Split-seq provides an affordable method for identifying cluster-specific isoforms in single cells that can be further quantified with companion deep short-read scRNA-seq from the same cell populations.

## Introduction

Alternative transcript isoform expression is a major regulatory process in eukaryotes that includes differential TSS (transcription start site) selection, RNA splicing, and TES (transcription end site) selection. These differential choices sculpt the transcriptome and its resulting proteome during development, across cell types and in disease states. However, it has proved challenging to fully capture and quantify isoform regulation by standard short-read RNA-seq because of the ambiguity it leaves in mapping the transcript termini and full-length exon connectivity that define each mature isoform.

In recent years, long-read RNA sequencing technologies have emerged as a powerful alternative for transcript-level identification and quantification by going beyond the level of exon-usage to simultaneously identify novel isoforms with alternative TSSs, TESs, and exon combinations. Furthermore, long-read RNA-seq has been adapted to single-cell sequencing using high-throughput microfluidics-based methods (1, 2, 3, 4). Some of these studies sequenced the same cells with both PacBio and Illumina technologies and relied on short-read gene quantification to cluster and characterize cell types, while using the long reads to identify full-length isoforms (2, 4). However, these prior approaches used expensive equipment, such as microfluidics platforms, and/or applied very high amounts of long-read sequencing whose expense limits routine and extensive application.

Differential RNA isoforms discriminate cell types within complex tissues and, within cell types such as neurons, can further distinguish functionally distinct cell sub-populations (5, 6). Isoform choice can even distinguish individual neurons of the same “type” from each other (7, 8). Transcript isoforms also discriminate developmental stages and disease states (9). In vertebrate systems, differential isoform regulation through development has long been appreciated, and in some disease states such as type 1 myotonic dystrophy, fetal or neonatal stage isoforms of *Tnnt2, Atp2a1* (*Serca1*), and *Ldb3* (*Zasp*) are inappropriately expressed (10, 11, 12). In addition, several studies have characterized the diversity of gene expression within the population of nuclei from myotubes (13, 14). This prior work on skeletal muscle provides known instances of isoform choices that we can use to benchmark new methods for transcriptome profiling, while at the same time posing unanswered questions that require single-cell or single-nucleus long-read data such as nuclear specialization within myotubes.

*In vitro* differentiation of the myogenic C2C12 cell line from proliferating, mononucleated myoblasts to multinucleated myotubes is a widely used model of myogenesis due to transcriptional and morphological similarities to the *in vivo* process (15). A subset of cells under differentiation promoting conditions remain mononucleated and are called MNCs (16, 17). In adult muscle tissue, satellite cells are mononucleated muscle stem cells that can be stimulated to proliferate and differentiate to drive muscle repair (18). Expression of the satellite cell marker gene *Pax7* decreases as satellite cells are activated into proliferating myoblasts, while expression of myogenic regulatory factors (MRFs) such as *Myod1* and *Myog* increase and promote myogenesis (18). Satellite cells undergo asymmetric divisions to produce future *Pax7* negative, MRF positive myoblasts and to self-renew *Pax7* positive, MRF-negative satellites (19). In addition to major transcriptional changes during myogenesis, C2C12 differentiation exhibits substantial changes, both qualitative and quantitative, in splice isoforms (20). For example, *Pkm* undergoes an isoform switch during C2C12 differentiation that results in two distinct isozymes of the gene, PMK2 and PKM1 (21). Proliferating C2C12s express both isoforms of beta-tropomyosin (*Tpm2*), including exon 6a or exon 6b, but expression of the 6b isoform increases substantially during differentiation (21).

Here, we combine combinatorial barcoding of individual C2C12 cells and nuclei using the Split-seq strategy (22) with long-read sequencing (LR-Split-seq) to investigate isoform changes during differentiation. We first examined the technical differences between LR-Split-seq random hexamer and oligo-dT priming strategies as well between single-cell and single-nucleus. We compared the performance of LR-Split-seq to bulk long-read RNA-seq, and further compared the clusters recovered from LR-Split-seq to those from short-read sequencing for the same cells, as well as a companion dataset of 37,000 cells to show that long-read single-cell transcriptomes produce similar results to short-read that can be readily integrated. We then leveraged LR-Split-seq results to identify and quantify TSSs in order to perform differential TSS testing and examine TSS usage between single-cell clusters. Finally, we integrated the resulting TSS expression from LR-Split-seq with matching single-cell ATAC-seq to quantify the extent of coordinated single-cell chromatin accessibility.

## Results

### Comparing oligo-dT versus random hexamer primed long-read data

Split-seq uses a combination of oligo-dT and random hexamer primers in order to decrease the 3’ bias that dominates other single cell RNA-seq methods that prime only with oligo-dT (22). These methods are designed to perform 3’ end counting for sequenced genes but they give little or no information about the rest of the transcript. In contrast, when Split-seq is conventionally performed with short reads, the random priming feature should, in the ideal instance, provide comprehensive information about the entire body of the transcript. However, this benefit in the short-read format is expected to impinge differently and not entirely favorably on long-read data. The extent and character of effects from internal priming will depend on multiple protocol variables (e.g. relative amounts of oligo-dT versus random hexamers, substrate RNA integrity) and on filtering steps in the subsequent informatic pipeline. We therefore began by testing the impact of priming strategy on the LR-Split-seq data. We collected proliferating C2C12 myoblasts (0hr) as both whole cells and nuclei, then differentiated the remainder into myotubes over 3 days to recover 72hr differentiated nuclei (Methods). We labeled a total of approximately 37,000 cells/nuclei from the three samples using the Split-seq combinatorial barcoding strategy. We then built a sub-library of 1,000 cells for sequencing by PacBio as well as Illumina (Fig. 1A). The LR-Split-seq data was first debarcoded and demultiplexed using our LR-splitpipe pipeline (Methods). We then analyzed the reads with TALON (23), which is designed to assign long reads to their transcripts of origin and to identify new transcripts (Fig. S1A-C) (Methods). TALON’s long-read RNA-seq annotation then assigns each read to a category that specifies whether the read matches a known transcript in the reference transcriptome GTF file, or if it represents a novel transcript (23, 24). Random hexamer priming is expected to start within the body of a transcript rather than the 3’ polyA tail where oligo-dT primers hybridize, though intronic A-rich runs are known to serve as additional start points for oligo-dT priming (25). This mixed priming strategy, as it is currently implemented in the Split-seq commercial platform, produced remarkably little difference in the final LR-Split-seq read length distribution from the two primer types (Fig. 1B, 1C). The distribution of reads per TALON category showed a slightly higher proportion of incomplete splice match (ISM) reads per cell from the random hexamer priming strategy versus the oligo-dT priming strategy (Fig. 1C). We speculate that the high fraction of oligo-dT primed reads per cell that begin at internal sites (∼60%) accounts for the overall similarity of random hexamer primed reads in length profiles and genes detected.

**Fig. 1:**
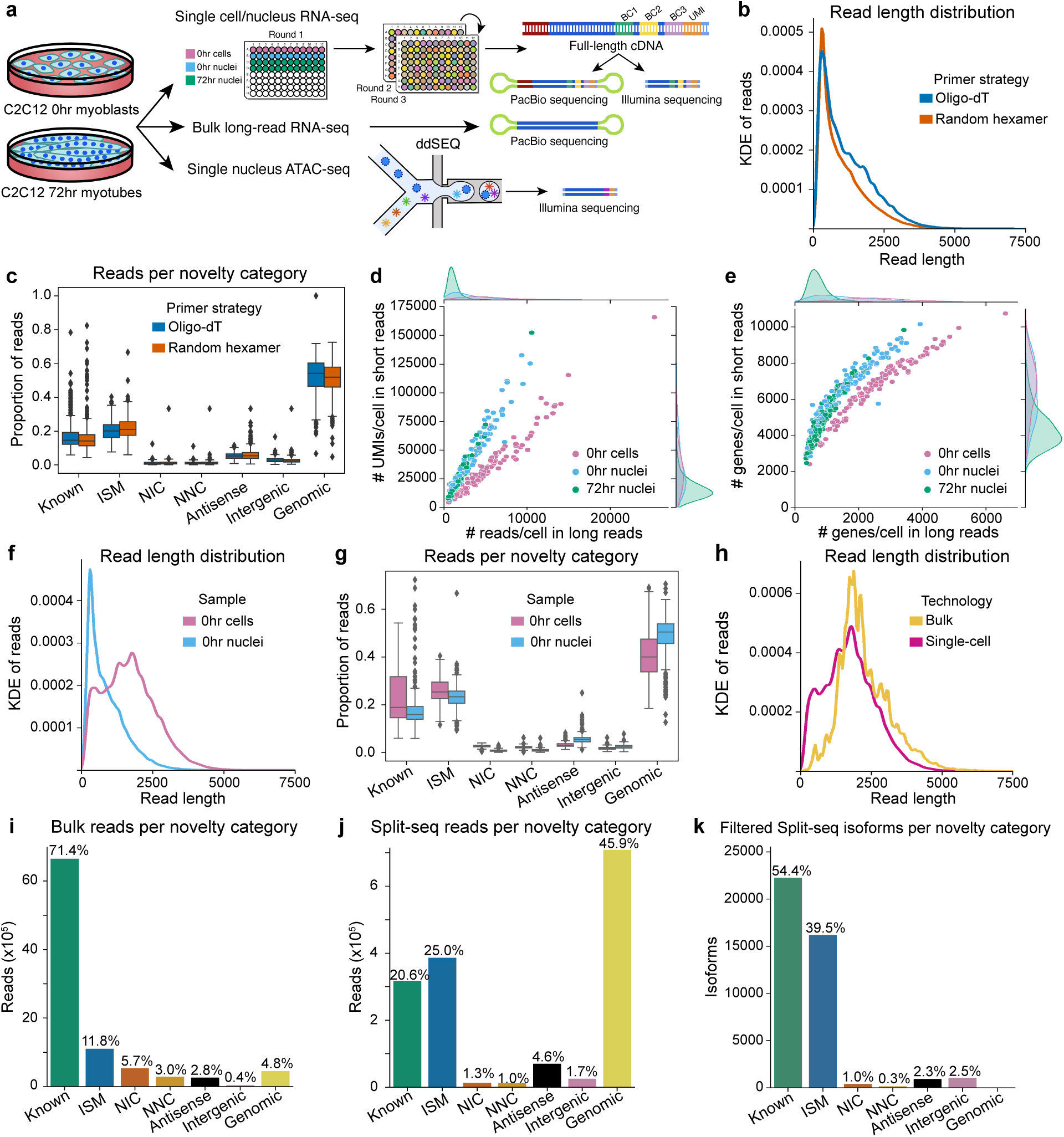
**Technical comparisons in LR-Split-seq and bulk long-read RNA-seq. a**, Schematic diagram of experimental design. Single cell/nucleus LR-Split-seq, bulk long-read RNA-seq, and single nucleus ATAC-seq were performed on C2C12 0hr myoblasts and 72hr differentiating cells. The same single-cell/UMI-barcoded cDNA was used in both short-read and long-read sequencing. **b,** Kernel density estimation (KDE) of read length distribution of oligo-dT primed reads (blue) compared to random hexamer primed reads (orange). **c**, Proportion of oligo-dT/random hexamer reads in each cell for each novelty category. **d**, Comparison of number of reads and **e**, genes detected between short and long reads. Cells are labeled by sample type (0hr cells in pink, 0hr nuclei in blue, and 72hr nuclei in green) and marginals on the top and right indicate their distributions. **f**, KDE read length distribution of 0hr cells (pink) compared to 0hr nuclei (blue) reads, not including genomic reads. **g**, Proportion of 0hr cell (pink)/nuclei (blue) reads per cell/nucleus per novelty category. **h**, KDE read length distribution of bulk long reads (yellow) compared to single-cell long reads (magenta), not including genomic reads. **i**, Unfiltered reads per novelty category in bulk long-read data and **j**, LR-Split-seq data. **k**, Filtered isoforms per novelty category across all cells in LR-Split-seq data.

### Single nuclei compared with single cells for LR-split-seq

We compared single-cell versus single-nucleus LR-split-seq. Overall, more reads and genes were recovered from whole cells versus nuclei for both long- and short-read data, which is expected because cytoplasmic transcripts are left behind during nuclear extraction, making the nuclear determinations less sensitive on a per cell basis (Fig. 1D, 1E). When comparing only 0hr cells with companion nuclei, we observe shorter read lengths in the nuclei (Fig. 1F). And as expected, we also see a larger proportion of genomic reads per cell/nucleus in nuclei compared to cells (Fig. 1G). These nuclear genomic reads could result from the enrichment of intronic RNA in the nucleus which would explain the lack of splice junctions.

Comparing LR-Split-seq of whole cells with bulk long-read RNA-seq for myoblasts, we found that the LR-Split-Seq is modestly shorter than bulk long-read data (Fig. 1H, Table S1). Bulk reads have an average mean length of 2,274 bp and a peak from the kernel density estimate (KDE) distribution of 1,875 bp, versus an average mean length of 1,735 bp and a KDE peak of 1,791 bp for LR-Split-seq non-genomic reads from whole cells (Fig. 1H, Table S1). The LR-Split-seq reads also had more genomic and incomplete splice match (ISM) reads than the bulk data (Fig. 1I, 1J). These differences are in line with expectations, given other differences in details of the bulk protocol (Methods). Nevertheless, after filtering our novel transcripts with TALON, the majority of transcript models we recover are annotated in GENCODE which we call known (Methods). This filtering resulted in 40,983 isoforms distributed across seven novelty categories (Fig. S1D, Fig. 1K). The observed read length differences between LR-Split-seq and bulk is reflected in the genes and transcripts that are uniquely detected in the bulk or LR-Split-seq. Transcripts detected only in bulk transcriptomes were likely to be longer, whereas transcripts detected only in LR-Split-seq data were enriched for shorter length (Fig. S1E, S1F). Due to overall longer read length in bulk long reads, these data were more likely to have multiple exons than LR-Split-seq (Fig. S1G). We conclude that the read length profile of known reads in single-cell LR-Split-seq is quite similar to bulk long reads, given protocol differences. This suggests to us that the overall shorter lengths in single-nucleus versus whole cell LR-split-seq are of biological origin, likely driven by underlying differences between cytosolic RNA, which is rich in mature mRNA versus nuclear RNA, which contains mature mRNA but in lower proportions.

### LR-Split-seq and bulk long-read RNA-seq detect similar gene sets

Despite differences in transcript length and novelty classification between bulk long-read RNA-seq and LR-Split-seq, we detected 9,584 known genes in both bulk and single-cell LR-Split-seq, with 5,195 of these shared across all assays and sample combinations (Fig. 2A). These results demonstrate the gene detection sensitivity of LR-Split-seq. The next largest intersections contain >1,500 genes recovered in all but the single-nucleus data which is likely due to the relative loss of cytoplasmic transcripts from the nuclear preparation. Genes detected in LR-Split-seq but not in the companion bulk RNA-seq tend to be short and are enriched for short RNA biotypes such as snoRNAs and miRNAs, while genes detected solely in bulk data are enriched for protein coding genes (Table S2). A plausible explanation is that Split-seq’s random hexamer priming captured these transcript types whereas the bulk method, which uses oligo-dT priming exclusively, preferentially captured polyadenylated transcripts. We also examined the overlap between filtered novel transcript models from the NIC and NNC novelty categories in bulk and LR-Split-seq. While the vast majority of novel transcript models were only reproducible between bulk replicates, 251 NIC transcripts and 61 NNC transcripts were reproducible in at least one bulk and one LR-Split-seq sample (Fig. S1H, S1I). These represent isoforms that are most likely to be real, though not previously annotated.

**Fig. 2:**
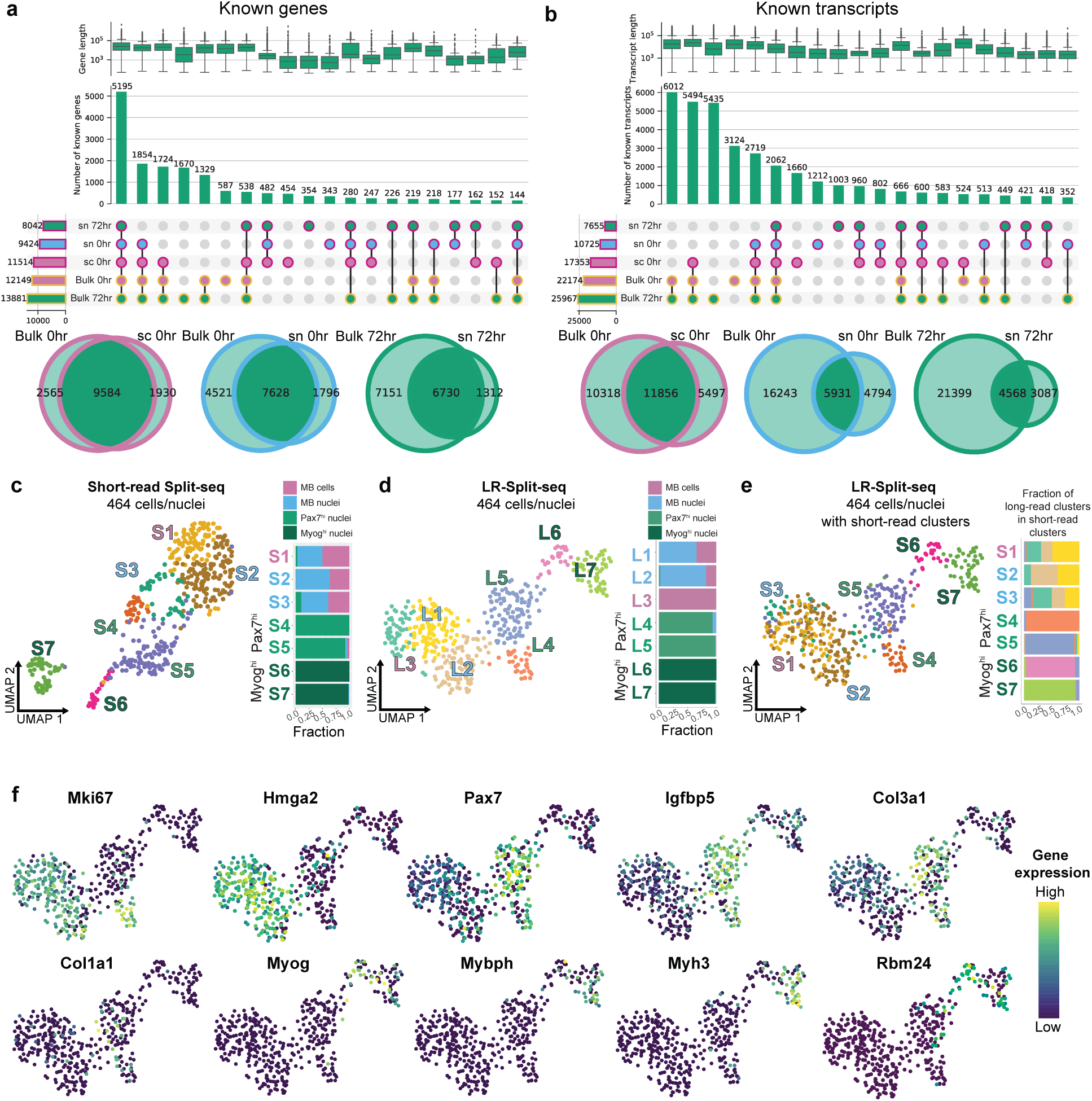
**LR-Split-seq in C2C12 0hr and 72hr samples recapitulates results from companion bulk and standard short-read Split-seq. a**, Upset plots of known genes found in bulk compared to LR-Split-seq data across all samples. Bars on the left indicate set size, circles indicate combinations of samples, and bars on top indicate the number of genes found in each combination (first 20 combinations shown). Outline colors indicate technology (bulk in yellow, single-cell or single-nucleus in magenta) and fill colors indicate sample type (72hr nuclei in green, 0hr nuclei in blue, and 0hr cells in pink for single-cell data; 72hr in green, 0hr in pink for bulk data). Box plots above indicate gene length distribution for each intersection. Venn diagrams below summarize the overlaps between bulk (left) and single-cell or single-nucleus (right), for each sample type. Sample type is indicated by outline color. **b**, Upset plot and Venn diagrams of known transcripts found in bulk data and LR-Split-seq data (first 20 combinations shown). **c**, UMAP of 464 short-read Split-seq cells/nuclei labeled by 7 Leiden clusters (S) and breakdown of cell type per cluster: 110 0hr cells (pink), 145 0hr nuclei (blue), and 209 72hr nuclei (*Pax7^hi^* in green and *Myog^hi^* in dark green). **d**, UMAP of 464 LR-Split-seq cells/nuclei using gene-level data labeled by 7 Leiden clusters (L) and **e**, Leiden cluster ID of matching short-read data (S) shown in **c**, as well as long-read cluster makeup of each short-read cluster. **f**, Expression of marker genes, dark blue = lowly expressed, yellow = highly expressed.

### LR-Split-seq recapitulates cell classifications recovered from short-read Split-seq

Overall, we recovered 110 0hr myoblast cells, 145 0hr myoblast nuclei, and 209 72hr differentiating nuclei (464 cells total) that passed short-read QC thresholds as well as an additional requirement of ≥500 long reads per cell in the 1,000-cell library (Methods) (Fig. S2A-E). Leiden clustering based on short-read sequencing of the 464 cells/nuclei yielded 7 clusters (S1-S7). We observed mixed populations of 0hr myoblast cells and nuclei in clusters S1-S3, while the 72hr differentiating nuclei clustered in S4-S7. This overall structure is consistent with differentiation playing a dominant role in the UMAP structure, while differences between nuclei versus whole cells from the 0hr samples were minor by comparison (Fig. 2C). Additional patterns in the dataset that agree with known biology in the system include expression of the satellite cell marker gene Pax7, which is expressed mainly in 72hr clusters S4 and S5, while the key myogenic transcription factor *Myog* (myogenin) is expressed mainly in 72hr clusters S6 and S7 (Fig. S2F). An independent Leiden clustering performed using the LR-Split-seq data for the same 464 cells proved very similar to the companion short-read clustering with 7 clusters (L1-L7) in which the myoblast progenitor cells/nuclei are in clusters L1-L3 while the differentiating sample gives rise to clusters L4-L7 (Fig. 2D). This UMAP again separates the latter group into *Pax7^hi^* (L4, L5) and contrasting *Myog^hi^* sets (L6, L7), with the latter expressing additional downstream markers of myocyte differentiation. Color-coding cells in the long-read UMAP according to the cluster identity from the companion short-read data showed high concordance of clusters L4-L7 with S4-S7 (Fig. 2E). The myoblast progenitor clusters S1-S3 and L1-L3 also agree, although the short-read clusters were more mixed between 0hr cells and nuclei. We investigated gene expression patterns for additional known marker genes across the cells and nuclei between the short-read and long-read clusters (Fig. 2F). Most notably, *Mybph*, *Myh3*, and *Mef2c* are highly expressed in a subset of 72hr nuclei that make up cluster L7, whereas *Myog* is expressed in both clusters L6 and L7 of 72hr nuclei (Fig. 2F, Fig. S2F). Similar to the short-read data, *Pax7* is present in both 0hr and 72hr clusters, but it is most highly expressed in clusters L4 and L5 (Fig. 2F). We also capture similar expression patterns in short-read and long-read *Pax7^hi^* 72hr subclusters as indicated by *Igfbp5*, *Col3a1*, and *Col1a1* (Fig. 2F, Fig. S2F). Due to the consistent expression patterns of known marker genes across both technologies, we postulate that *Myog^hi^* clusters S6, S7, L6, and L7 are mainly nuclei originating from fused, multinucleated myotubes or mononucleated myocytes on their way toward fusion, while the *Pax7****^hi^*** clusters S4, S5, L4, and L5 are nuclei distinct from both myoblasts and the 72hr *Myog^hi^* nuclei.

To harness the full-length capacity of LR-Split-seq, we performed isoform switching tests with a corrected p-value cutoff from a chi-squared test of 0.05 and a change in percent isoform usage cutoff of ≥10% (4) (Methods). We performed this pairwise test across three identified groups of clusters: 0hr myoblast (MB) cells (L1-L3), 72hr *Pax7****^hi^*** nuclei (L4-L5), and 72hr *Myog^hi^* nuclei (L6-L7). We recovered statistically significant isoform switching genes that have been previously observed in differentiating C2C12s, such as *Tpm2* (*Adj. P* = 2.02×10^-5^ MB vs. 72hr *Myog^hi^*) and *Pkm* (*Adj. P* = 4.75×10^-11^ MB vs. 72hr *Pax7****^hi^***; *Adj. P* = 5.69×10^-7^ MB vs. 72hr *Myog^hi^*). The *Tpm2* locus specifically shows an increase in expression of and preference for isoforms containing exon 6b in the differentiated nuclei as previously characterized in C2C12s as visualized with Swan (Fig. S2G) (21, 26). We found 21 significant isoform-switching genes between MB nuclei and 72hr *Pax7****^hi^*** nuclei as well as 13 significant isoform-switching genes between MB nuclei and 72hr *Myog^hi^* nuclei (Table S3, S4).

### C2C12s have distinct *Pax7^hi^* subpopulations following differentiation

We confirmed the presence of distinct *Pax7^hi^* and *Myog^hi^* clusters by short-read sequencing of an extended set of cells and nuclei from the same labeled pool, comprised of six additional 9,000-cell sub-libraries on top of the 1,000-cell sub-library with matching long reads (Methods). After filtering, we recovered 36,869 total cells/nuclei from all seven sub-libraries, including the 464 cells/nuclei with both short and long reads (Fig. S2A-E). The 7,797 myoblast cells, 10,194 myoblast nuclei, and the 18,878 differentiating condition nuclei clustered primarily by differentiation state (Fig. 3A). The progenitor states formed one main group in UMAP space that slightly separates cells and nuclei, while the differentiating nuclei extend outward in a spectrum with several smaller groups (Fig. 3A). Of the 20 clusters identified by Leiden clustering, 7 consist mostly of myoblast cells/nuclei while 13 are mainly differentiating nuclei (Fig. 3A) (Methods). Out of the 13 72hr clusters, 8 are *Pax7^h^*^i^ and the other 5 are *Myog^hi^*, which is consistent with results from the 464 cells alone (Fig. 3A). Accordingly, cells from each of the 20 clusters are represented by both short and long reads in the 464- cell subset (Fig. 3B, 3C). We assign these clusters to the cells we recovered with long reads to better inform the cellular identities with high resolution (Fig. S2H). For example, a small subset of 12 cells out of 105 total cells in cluster S5 belong to cluster R12, which is distinguished by high expression of *Col1a1* (Fig. 3D). Genes critical for cell cycle phases G1 and S such as *Cdk2* and *Pcna* are highly expressed in MB cluster R1, while G2 and M phase marker gene *Top2a* is highly expressed in MB clusters R2 (made up of mostly cells) and R3 (made up of mostly nuclei) as well as *Pax7^hi^* cluster R9 (Fig. S2I) (27, 28, 29). *Myog* and myogenic marker gene *Mybph* are highly expressed in clusters R16, R17, R18, and R20, indicating that these nuclei most likely belong to committed myocytes and myotubes (Fig. 3D). RNA velocity analysis, which uses the ratio of intronic (unspliced) and exonic (spliced) reads to predict the transcriptional trajectory of cells, reveals a lineage from clusters R17 and R18 toward clusters R19 and R20. R19 and R20 express terminal myogenic marker genes such as *Myh3*, *Mef2c*, *Tnnt2*, and *Neb* (Fig. 3D, Fig. S3A) (Methods). Of the 8 co-adjacent *Pax7^hi^* clusters (R8, R9, R10, R11, R12, R13, R14, and R15), some also express cluster-discriminating genes such as *Igfbp5* (cluster R11), *Col1a1* (cluster R12), and *Itm2a* (cluster R14) (Fig. 3D, Fig. S3A). We validated differential cluster-specificity of marker genes using spatial transcriptomic profiling of *Col1a1* (cluster R12), *Itm2a* (cluster R14) and *Myh3* (cluster R20), which showed patterns fully consistent with the Split-seq data (Fig. 3E). Imaging also confirmed that *Pax7^hi^* subcluster marker genes are expressed in MNCs rather than in the multinucleated myotubes that they surround (Fig. 3E). *Myh3* is expressed throughout multinucleated myotubes but less so in mononucleated cells. *Pax7^hi^* MNCs appear to either express *Col1a1* or *Itm2a,* consistent with their mutually exclusive marking of clusters R12 and R14 (Fig. 3D, 3E). *Myog* is expressed throughout multinucleated myotubes as well as in some mononucleated cells that are likely to be pre-fusion myocytes (Fig. S3B).

**Fig. 3:**
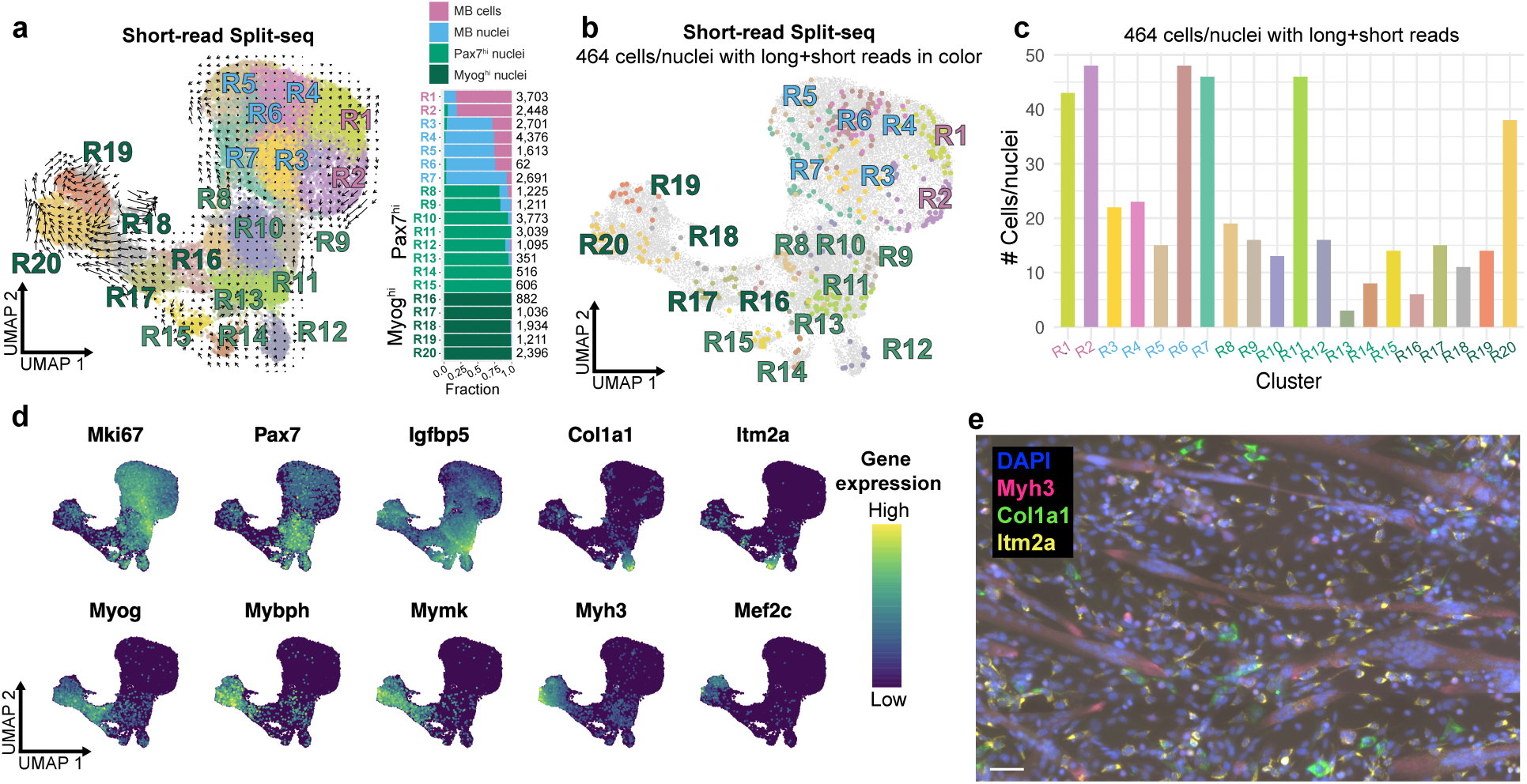
**Short-read Split-seq analysis. a**, UMAP of 36,869 short-read Split-seq cells/nuclei labeled by 20 Leiden clusters (R) with RNA velocity field trajectories and breakdown of cell type per cluster with number of cells per cluster: 7,797 0hr myoblast cells (pink), 10,194 0hr myoblast nuclei (blue), 18,878 72hr nuclei (*Pax7^hi^* in green and *Myog^hi^* in dark green). **b**, UMAP of short-read Split-seq cells/nuclei with the 464 cells with matching long reads in color corresponding to R1-R20. **c**, Histogram of the number of the 464 cells/nuclei per R1-R20. **d**, Distribution of expression of marker genes; dark blue = lowly expressed, yellow = highly expressed. **e**, Visualization of transcripts in mononucleated cells and myotubes at the 72hr differentiation timepoint. Blue = DAPI, pink = *Myh3*, green = *Col1a1*, yellow = *Itm2a*. Scale bar: 50 μm.

We observed heterogeneous populations of differentiating cells representing cell populations and states that are involved in adult muscle tissue repair. Clusters R10 and R11 express *Igfbp5*, which promotes muscle differentiation, and *Nfix*, which controls timing of regeneration by repressing myostatin (Fig. 3D, Fig. S3C, Table S5) (30, 31). Cluster R12, marked by *Col1a1*, *Fn1* (fibronectin), and a number of other collagen genes, may represent a population of previously defined MNCs that can transiently remodel their ECM, which is a process shown to regulate satellite cell numbers *in vivo* (Fig. 3D, Fig. S3A, Table S5) (32, 33). Cluster R13 expresses *Lix1*, a *Pax7* target gene needed for activated satellite cell proliferation (Fig. S3A, Table S5) (34). Cluster R14, which expresses *Itm2a* and *Pax7,* may be analogous to activated satellite cells (Fig. 3D, Fig. S3A, Table S5) (35). Appropriately, the cluster R14 RNA trajectory tends toward cluster R15 which expresses *Tead1* (*Tef-1*) and *Myog,* which are known to promote muscle differentiation (Fig. S3C, Table S5) (36).

### Chromatin accessibility of myogenic marker genes distinguishes *Myog^hi^* and *Pax7^hi^* 72hr nuclei

To assess chromatin accessibility in the groups of nuclei we identified with LR-Split-seq, we performed snATAC-seq on matching timepoints. We recovered 23,525 single nuclei from our snATAC-seq experiments following filtering and QC (Fig. S4A-B), resulting in 18 clusters from Leiden clustering: seven 0hr myoblast clusters and eleven 72hr differentiating clusters (Fig. 4A) (Methods). Gene activity is a measure of chromatin accessibility of the gene body and 2kb upstream as a rough estimate of transcriptional activity (37). We saw that chromatin gene activity patterns in our snATAC-seq UMAP for *Myog* is somewhat similar to scRNA-seq expression patterns, where the *Myog* locus was highly accessible in a subset of differentiated clusters (A16, A17, and A18) (Fig. S4C). To investigate the agreement between expression and chromatin accessibility for the same time points, 0hr and 72hr, we integrated our short-read Split-seq and snATAC single-cell measurements using Signac (Methods) (38). This integration mapped Split-seq cells on snATAC-seq nuclei, resulting in predicted snATAC-seq cell types. The predicted Split-seq time point (0hr or 72hr) was mostly accurate, with 96% (10,136 out of 10,508) of true snATAC 0hr nuclei predicted to be 0hr from the expression data and 79% (10,381 out of 13,017) of true snATAC 72hr nuclei predicted as 72hr (Fig. S4D). When we mapped Split-seq cells grouped by MB (R1-R7), *Pax7^hi^* (R8-R15), and *Myog^hi^* (R16-R20) onto snATAC nuclei, we found that 48% (1,502 out of 3,148) of nuclei with a *Myog* activity score > 0 were predicted to be *Myog^hi^* and that 27% (5,135 out of 18,542) of nuclei with a *Pax7* activity score > 0 were predicted to be *Pax7^hi^* (Fig. S4D). Unlike our Split-seq RNA data, where we detected high expression of *Pax7* in specific clusters, ATAC-based gene activity scores predicted that *Pax7* would be equally active across all clusters (Fig. S4E). Taken at face value, this suggests that some differentially expressed genes do not exhibit corresponding changes in promoter chromatin state, as reflected by these activity scores. However there are several distal peaks ATAC peaks located downstream of *Pax7* whose dynamics are coordinated with the RNA. This suggests, as a working model, that they are regulatory elements governing *Pax7* expression. In contrast, *Myog* and *Mybph* illustrate expected coordinated changes in chromatin accessibility and RNA isoform expression during differentiation (clusters A16-A18) at the TSSs of these genes (Fig. S4F). For uniform terminology between RNA and DNA data, we label 72hr *Myog^low^* snATAC clusters A8-A15 as *Pax7^hi^*. While snATAC can clearly capture changes in chromatin remodeling, the ATAC-only gene activity scores (at least as computed by Signac) do not reflect the *Pax7* expression level changes that we measure in this system.

**Fig. 4:**
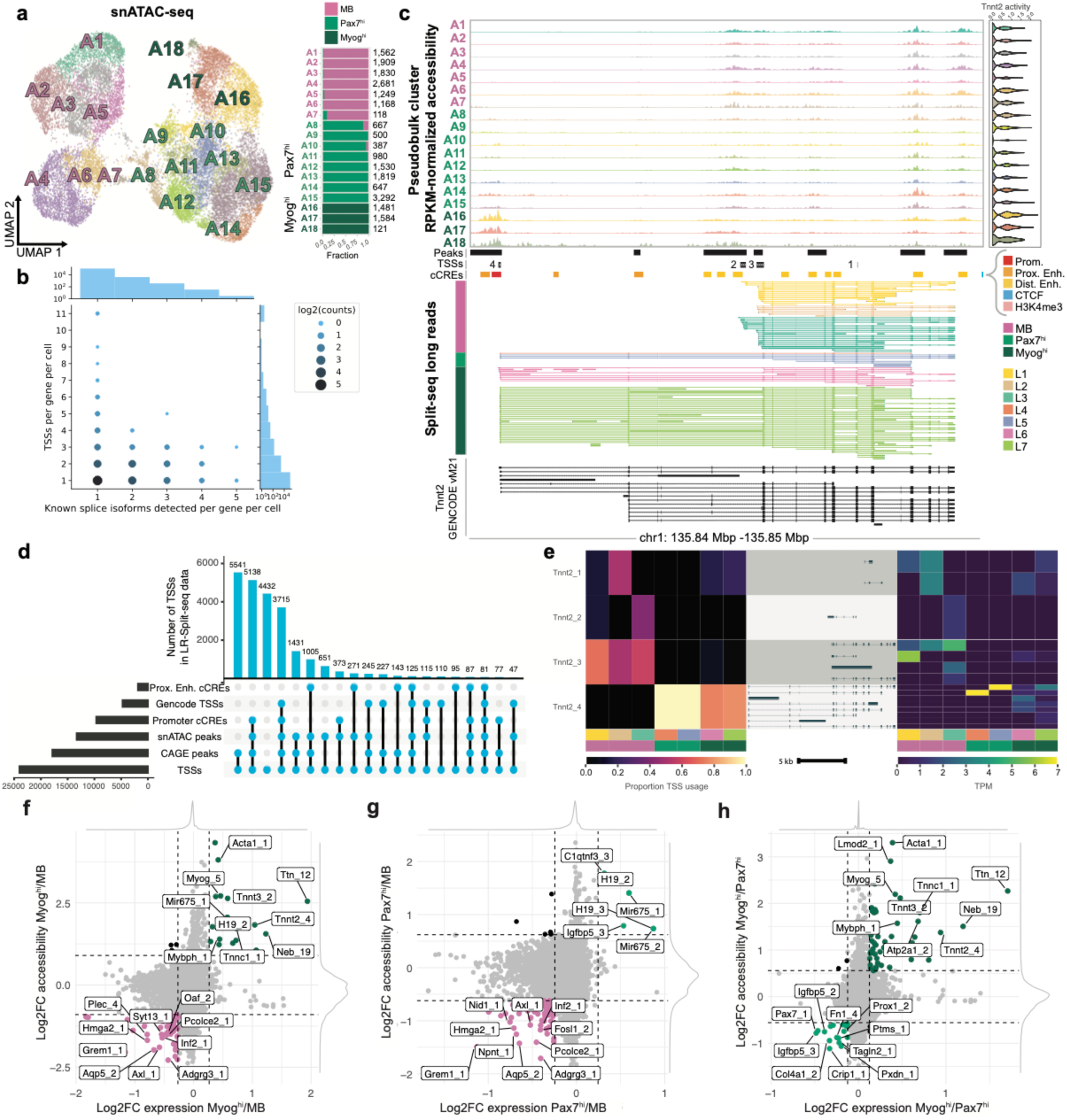
**Identification of TSSs from LR-Split-seq and integration with snATAC-seq. a**, UMAP of 23,525 snATAC-seq nuclei labeled by 18 Leiden clusters (A) and breakdown of cell type per cluster with number of cells per cluster on right: 10,508 0hr myoblast nuclei (pink) and 13,017 72hr nuclei (*Pax7^hi^* in green and *Myog^hi^* in dark green). **b**, Bubble plot of the number of distinct known splice isoforms per gene per cell compared to the number of distinct TSSs per gene per cell in LR-Split-seq. **c**, Track plot of alternative *Tnnt2* TSS usage between 72hr differentiating cells and 0hr myoblasts. From top to bottom: clustered snATAC-seq pseudobulk peaks, merged psuedobulk peaks, TSS regions called from LR-Split-seq, ENCODE cCREs, clustered LR-Split-seq reads used to call TSSs, and comprehensive set of GENCODE vM21. **d**, Validation of TSSs found in LR-Split-seq using four external datasets and snATAC-seq pseudobulk peaks (first 20 intersections shown). **e**, Left, proportion of TSS-assigned reads in LR-Split-seq clusters from each identified *Tnnt2* TSSs. Right, expression of each TALON filtered *Tnnt2* isoform in LR-Split-seq clusters with corresponding transcript models associated with each *Tnnt2* TSS. **f**, Comparison of log2 fold change (LFC) in expression and accessibility across identified TSSs: *Myog^h^*^i^ (+LFC) compared to MB (-LFC), **g**, *Pax7^hi^* (+LFC) compared to MB (-LFC), and **h**, *Myog^hi^* (+LFC) compared to *Pax7^hi^* (- LFC).

As expected, investigation of marker peaks for *Myog^hi^* clusters A16-A18, using a gene annotation method with gene ontology analysis, revealed significant terms such as muscle system process (*P* = 1.55 x 10^-115^), muscle structure development (*P* = 5.77 x 10^-118^), and striated muscle contraction (*P* = 3.87 x 10^-96^) (Methods, Fig. S3G, Table S6, Table S7). In comparison, MB clusters A1-A7 had broad significant terms such as regulation of anatomical structure morphogenesis (*P* = 1.69 x 10^-19^), cell-cell adhesion (*P* = 3.45 x 10^-13^), and cell motility (*P* = 3.89 x 10^-14^) (Table S6, Table S7). The significant terms for *Pax7^hi^* clusters A8-A15, in contrast to *Myog^hi^* clusters, were extracellular matrix organization (*P* = 1.23 x 10^-9^), extracellular structure organization (*P* = 1.35 x 10^-9^), and blood vessel morphogenesis (*P* = 3.92 x 10^-9^) (Table S6, Table S7). Most marker peaks defining the *Myog^hi^* clusters are specific to skeletal muscle myogenesis in myotubes while marker peaks for *Pax7^hi^* clusters indicate that they have a supportive role during development, such as by providing structural integrity to myotubes through ECM remodelling.

### LR-Split-seq identifies differential TSS choice

We developed a peak calling script to identify TSSs and TESs from long-read data (Methods). For both bulk and single-cell data, reads filtered by known, NIC, NNC, and prefix ISMs for TSSs or suffix ISMs for TESs were scanned with a window of 50bp to call TSS and TES peaks. Each end was required to be supported by at least 2 long reads (Fig. S5A). We further filtered the ends at the level of each gene to achieve a refined set of TSSs and TESs for the bulk and LR-Split-seq data separately: 22,938 TSSs in bulk (Fig. S5B, S5C), 23,996 TSSs in LR-Split-seq (Fig. 4B), 14,120 TESs in bulk (Fig. S5D, S5E), and 12,521 TESs in LR-Split-seq (Fig. S5F, S5G, Methods, Tables S8-11). Comparing the number of distinct ends to the number of distinct splice isoforms revealed that multiple TSSs are expressed per single splice isoform in both bulk and single cells (Fig. 4B, S5B). *Tnnt2* (troponin T2) has multiple known isoforms (39) and is differentially expressed between *Myog^hi^* and *Pax7^hi^* nuclei in the short-read data, so we decided to investigate chromatin accessibility and TSS usage at the *Tnnt2* locus (Fig. S3A, Table S5, Fig. 4C). We recovered four distinct TSSs for *Tnnt2*, three of which (Tnnt2_2, Tnnt2_3, and Tnnt2_4) overlap snATAC pseudobulk peaks, and all four of which overlap prior CAGE peaks found in C2C12 (40). Tnnt2_4 overlaps a known promoter cCRE and GENCODE vM21 transcript start site, while Tnnt2_2 overlaps a distal enhancer cCRE (Fig. 4C) (41). Tnnt2_4 has both higher expression in the LR-Split-seq data and increased accessibility in snATAC *Myog^hi^* and *Pax7^hi^* clusters, while Tnnt2_2 and Tnnt2_3 are more highly expressed and accessible in MB clusters (Fig. S5H-J). Therefore, an isoform switch occurs in *Tnnt2* where *Myog^hi^* and *Pax7^hi^* nuclei mainly use the known TSS belonging to the longer isoform, while the MB nuclei mainly use TSSs belonging to shorter isoforms. Genome-wide, we validated our TSS calls using an extended set of data: our snATAC pseudobulk peaks, GENCODE vM21 TSSs, ENCODE cCREs (promoter and proximal enhancer) from mm10, and C2C12 CAGE peaks, and found that the majority of the TSSs identified from LR-Split-seq are validated by at least one of these five other datasets (Methods, Fig. 4D).

Using the same strategy we implemented to detect isoform switching genes, we performed differential TSS usage tests on our LR-Split-seq data (Methods). We again subset our LR-Split-seq data into 0hr MB nuclei, 72hr *Pax7^hi^* nuclei, and 72hr *Myog^hi^* nuclei, and performed pairwise tests. In the MB vs. *Pax7^hi^* comparison, we found 39 genes with differential TSS usage (Table S12). In the MB vs. *Myog^hi^* comparison, we found 40 genes with differential TSS usage. Consistent with our previous findings, this list includes *Tnnt2*, where the MB nuclei only express isoforms consistent with downstream TSSs (Tnnt2_1, Tnnt2_2, Tnnt2_3) (Table S13). Conversely, the *Myog^hi^* subset predominantly expresses isoforms using the upstream TSS (Tnnt2_4) (Fig. 4C, 4E).

Similarly, we found multiple distinct TESs per splice isoform in bulk and LR-Split-seq data (Fig. S5D, S5F). We validated genome-wide TESs using GENCODE vM21 TESs and polyA-seq peaks from C2C12 cells at days 0 and 4 of differentiation, which overlapped the majority of TESs found in bulk data but not in the LR-Split-seq data (Fig. S5E, S5G) (Methods).

### Coordination of chromatin accessibility with transcriptional output

We calculated snATAC TSS chromatin accessibility across our refined set of TSSs to determine which TSSs are supported by both differential accessibility and expression (Methods). We compared the average log2 fold change (LFC) in both accessibility and expression between *Myog^hi^* and MB (Fig. 4F), *Pax7^hi^* and MB (Fig. 4G), and *Myog^hi^* and *Pax7^hi^* (Fig. 4H). Between MB and *Myog^hi^*, 19 TSSs are specific to *Myog^hi^* with an average LFC greater than two standard deviations (indicated by dashed lines) in both datasets, and 70 TSSs are specific to MB with average LFC less than two standard deviations in both datasets (Fig. 4F, Table S14). Several of the genes with such TSSs are differentially expressed (Table S5, Table S14). Only 6 TSSs were *Pax7^hi^*-specific relative to MB, but one of these is *Igfbp5*, which is a gene that was highly differentially expressed in the *Pax7^hi^* subset (Fig. 4G, Fig. S3A,Table S3, Table S14). Comparing MB and *Pax7^hi^*, 77 TSSs are MB-specific, 36 of which are also MB-specific when comparing *Myog^hi^* with MB. Of the 19 *Myog^hi^*-specific TSSs between *Myog^hi^* and MB, 15 were also *Myog^hi^*-specific when compared to *Pax7^hi^* (out of 53 total) (Fig. 4H, Table S14). Several of the 17 *Pax7^hi^*-specific TSSs (Fig. 4H) belong to differentially expressed genes, such as *Pax7*, *Col4a1*, *Fn1*, and *Igfbp5* (Fig. S3A, Table S5, Table S14). From a biological perspective, *Prox1* and *Vgll4* are potentially interesting; although they were not differentially expressed in the short-read data, they are known to be involved in skeletal muscle regeneration (Fig. 4H, Table S14) (42, 43).

## Discussion

The first goal of this work was to advance our capacity to directly map and quantify RNA isoforms in single cells. Using the C2C12 myogenic differentiation as a test system, we introduce long read-Split-seq (LR-Split-seq) and show that it can be as effective as standard short-read Split-seq for detecting cell clusters, based on data from the same number of cells or nuclei. This conclusion applied to nuclei as well as whole cells, although whole-cell data detected more genes per cell than companion LR-Split-seq data from nuclei. For biological systems that do not permit uniform whole-cell disaggregation such as our multinucleated myotubes or brain tissue, the success shown here for nuclei is encouraging. We speculate that the remaining sensitivity differential between nuclei and whole cells is a consequence of the smaller starting number of transcripts in nuclei, and some of that could be further compensated by increasing the nuclear number sequenced and their depth of sequencing. We also suggest that combining random hexamer primed long-reads with the oligo-dT primed long-read data helped to capture 5’ ends that are critical for inferring TSS use, although this adds incomplete PacBio reads to the overall dataset. We also illustrate that LR-Split-seq affords users the choice of analyzing the oligo-dT primed and hexamer primed read populations separately. A second motive for developing LR-Split-seq is that it will allow flexible study designs that can efficiently and more economically refine cell type identities by integrating additional standard short-read Split-seq data on the same samples. Results presented here showed that this strategy was effective in refining stem cell identities and states in the C2C12 system. Finally, we integrate results from LR-Split-seq with snATAC to gain insights into the dynamics of chromatin accessibility at the corresponding promoters with a longer term goal of building a fully integrated model of physically or genetically affiliated distal regulatory elements.

We were able to detect 79% of the genes and 53% of transcript isoforms detected in bulk myoblast long-read RNA-seq using LR-Split-seq in single cells. We expect these differences relative to bulk samples to be a function of the individual study design, including number and diversity of cells sequenced, depth of sequencing, fixation protocol and, for isoform detection, the contribution from internal hexamer priming. The largest sets of genes detected across the entire analysis included the LR-Split-seq assays, supporting the conclusion that it detects expressed genes reliably and reproducibly. The differences between known gene and transcript detection rates, relative to bulk data, were largely attributable to internally-primed Split-seq reads and their management in our computation pipeline. Specifically, we used TALON, which leverages non-full-length reads for quantification and detection on the gene-level but not on the transcript-level. Consequently, we achieved high gene detection concordance but lower transcript detection concordance between long-read bulk and LR-Split-seq data.

Gene-level clusters in LR-Split-seq are remarkably similar to the results in the equivalent standard short-read Split-seq. In both assays, clusters of differentiating cells were most homologous to each other and were distinct from the myoblast clusters. However, in LR-Split-seq, there was a greater tendency for the clusters to separate by assay format, as shown in the 0hr myoblast cells and nuclei. We captured expression dynamics of well-known myogenic marker genes in the differentiating clusters such as *Pax7, Myog, Mybph,* and *Myh3* that are reproducible in the short-read data we sequenced from the same cells (18). The additional context from ∼37,000 short-read single cells allowed us to investigate the *Myog^hi^* clusters in greater detail. We found that *Myog^hi^* clusters were very distinct from MB clusters, while *Pax7^hi^* clusters were in a spectrum of differentiation stages between MB and *Myog^hi^* clusters. Expression of additional marker genes in *Pax7^hi^* subclusters, RNA velocity trajectories, and validation with spatial transcriptomic profiling confirmed that these nuclei are from mononucleated cells in varying stages of differentiation.

LR-Split-seq enabled us to investigate transcript-level differences between the various stages of differentiation in myogenesis. We found novel insights into the biology of the system by studying differential TSS usage and integrating our TSSs identified from long reads and our snATAC-seq peaks. Our analysis revealed over 50 significant switches in TSS usage across clusters of undifferentiated versus differentiated stages, including a pronounced switch in *Tnnt2*, where the myoblasts primarily use TSSs that are novel to more recent GENCODE transcript annotations, while differentiated cells mainly express the known TSS that results in a longer isoform. This TSS switch was complemented by a corresponding increase in chromatin accessibility at the newly-expressed TSS in *Myog^hi^* clusters.

Unlike previous long-read scRNA-seq methods that rely on sequencing of each cell using custom microfluidics equipment (2, 4), LR-Split-seq is immediately accessible with no cell/droplet handling instrumentation and it is tunable in both cell number and sequencing depth, depending on the complexity of the underlying sample’s cellular composition. Additionally, it can be scaled up for long-read sequencing with additional sub-libraries and higher read depth. We believe that this technology and study design allow one to optimize the amount and character of information from short and long-read single-cell technologies when the costs of input cells, overall platform, and sequencing are all considered. While short-read Split-seq provides a broad survey of the transcriptional complexity of a biological system by sequencing up to 100,000 cells, corresponding LR-Split-seq can be applied to a targeted number of cells to provide higher-resolution isoform-level insights using a few million long reads from a few PacBio runs. In this way, LR-Split-seq promises affordable, simultaneous transcriptional profiling of a wide variety of tissues using short and long-read sequencing.

## Data / code availability

Data
| Description | GEO accession | ENCODE Accession |
| --- | --- | --- |
| Aggregate raw/processed short/long Split-seq data | GSE168776 | --- |
| Aggregate raw/processed snATAC-seq data | GSE168776 | --- |
| Bulk 0hr C2C12 biosamples | --- | ENCBS824FPY,<br>ENCBS649CMC |
| Bulk 72hr C2C12 biosamples | --- | ENCBS373BHL,<br>ENCBS606QKU |
| Split-seq parent C2C12 biosample | --- | ENCBS570IED |
| Split-seq 0hr C2C12 cells biosample | --- | ENCBS521YWL |
| Split-seq 0hr C2C12 nuclei biosample | --- | ENCBS431NOZ |
| Split-seq 72hr C2C12 nuclei biosample | --- | ENCBS978ZNQ |
| snATAC-seq 0hr biosample | --- | ENCBS081AJF,<br>ENCBS562OEW |
| snATAC-seq 72hr biosample | --- | ENCBS779SXF,<br>ENCBS143VME |
| snATAC-seq 72hr biosample (filtered) | --- | ENCBS247OBN,<br>ENCBS090IYH |

Code
| Code description | Link |
| --- | --- |
| Custom LR-Split-seq demultiplexer | https://github.com/fairliereese/LR-splitpipe |
| Figure generation code | https://github.com/fairliereese/2021_c2c12/ |

Additional resources
| Description | Link |
| --- | --- |
| Track hub for pseudobulk snATAC peaks per cluster, LR-Split-seq reads used for TSS calling per cluster, ENCODE cCREs, GENCODE vM21 transcript models | https://github.com/erebboah/c2c12_trackhub |

## Supplementary Tables

Table S1: LR-Split-seq TALON read annotation.

Table S2: Bulk vs. single-cell long read gene type detection test results.

Table S3: LR-Split-seq differential isoform test (MB vs. 72hr *Pax7^hi^*).

Table S4: LR-Split-seq differential isoform test (MB vs. 72hr *Myog^hi^*).

Table S5: Short-read Split-seq marker genes.

Table S6: snATAC-seq marker peaks.

Table S7: snATAC-seq GREAT results.

Table S8: LR-Split-seq TSSs

Table S9: LR-Split-seq TESs

Table S10: Bulk long-read TSSs.

Table S11: Bulk long-read TESs.

Table S12: LR-Split-seq differential TSS test (MB vs. 72hr *Pax7^hi^*).

Table S13: LR-Split-seq differential TSS test (MB vs. 72hr *Myog^hi^*).

Table S14: Log2FC TSS expression and accessibility.

## Methods

### C2C12 culture and differentiation

C2C12 cells from the American Type Culture Collection (ATCC, CRL-1772) were cultured on 10 cm plates (Thermo Scientific, 172931) in 10 mL myoblast growth media: high-glucose DMEM with L-glutamine and without sodium pyruvate (HyClone, SH30022.FS), supplemented with 20% fetal bovine serum (Omega Scientific, FB-11), 100 units/mL penicillin, and 100 ug/mL streptomycin (Gibco, 15140122). Cells were maintained at 20-50% confluency at 37°C with 5% CO_2_ and passaged at 1:3 or 1:4 every 2 to 3 days. All cells used in experiments were passaged less than 10 times. To detach them from plates, cells were rinsed with 1X PBS (HyClone, SH30256.02) and incubated with 2 mL TrypLE-Express (Gibco, 12605010) for 5 minutes at 37°C, which was then neutralized with 8 mL myoblast growth media. To differentiate, cells at 90-100% confluency were rinsed with 1X PBS and myoblast growth media was replaced with 10 mL differentiation media: high-glucose DMEM with L-glutamine and without sodium pyruvate (HyClone, SH30022.FS), supplemented with 2% donor horse serum (Gibco, 16050130), 100 units/mL penicillin, 100 ug/mL streptomycin (Gibco, 15140122), and freshly-added 1uM insulin (Sigma-Aldrich I6634). Differentiation media was replaced every 24 hours for 3 days. Cells were monitored under a microscope (EVOS FL Auto 2) to observe changes in morphology and confirm differentiation.

### Preparation of myoblast and myotube single-nucleus suspensions

We followed the Bio-Rad SureCell WTA 3’ Library Prep protocol for preparation of nuclei samples (44). Myoblasts from one 10 cm plate (∼1.5 million cells) and myotubes from one 10 cm plate (∼5 million cells) with >90% viability were lifted as described above and pelleted in 15 mL polypropylene falcon tubes (VWR, 89039-670) by centrifuging for 5 minutes at 1500 RPM. Cells were washed twice with cold 1X PBS + 0.1% BSA (Sigma-Aldrich A9418) and 0hr myoblasts were filtered through a 40 μm strainer; due to their size, 72hr samples containing myotubes were not filtered. After centrifuging for 3 minutes at 300 x g, cells were resuspended in 1 mL cold lysis buffer: 10 mM Tris-HCl pH 8 (Thermo Scientific, AM9855G), 10 mM NaCl (Fisher Scientific, S271), 3 mM MgCl_2_ (Sigma, M8266), 0.1% IGEPAL CA-630 (Thermo Scientific, 28324), 0.2 U/μL SUPERase In RNase Inhibitor (Invitrogen, AM2694) and 10 mg/mL BSA in nuclease-free water (Ambion, AM9937). Cells were incubated in lysis buffer on ice for 10 minutes, centrifuged at 4°C for 3 minutes at 300 x g, and washed with 1 ml of cold 1X PBS + 1% DEPC water (Invitrogen, 750023). The lysis, spin, and wash steps were repeated two more times for the 72hr samples because myotube cell membranes are more difficult to fully lyse than mononucleated myoblasts. Nuclei were stained with Trypan Blue (Bio-Rad, 1450021), and cell membrane lysis was confirmed under a microscope and by percent viability (<10%). Nuclei were stored on ice in 1 mL nuclei storage buffer (lysis buffer without the addition of IGEPAL CA-630).

### Preparation of single-cell barcoded cDNA using Split-seq

Single-cell barcoded cDNA and Illumina libraries were prepared using the Fixation Kit for Cells, Fixation Kit for Nuclei, and Single Cell Whole Transcriptome Kit (Parse Biosciences, SB2001) following the manufacturer’s protocols. Nuclei from the 0hr myoblast sample and 72hr sample in single-nucleus suspensions were counted on a TC20 Automated Cell Counter (Bio-Rad, 1450102), and ∼4 million were filtered through a 40 μm strainer into 15 mL polypropylene falcon tubes. Nuclei were fixed for 10 minutes and permeabilized for 3 minutes on ice, then DMSO was added for storage overnight at -80°C in a Mr. Frosty. Myoblast cells were similarly counted and filtered through a 40 μm strainer, followed by fixation and permeabilization. DMSO was added and cells were stored overnight at -80°C in a Mr. Frosty. Before storage, single-cell and single-nucleus suspensions were confirmed under a microscope.

To prepare barcoded cDNA, fixed and frozen cells and nuclei were thawed in a 37°C water bath and counted. Cells were added to the Round 1 reverse transcription barcoding plate at around ∼15,000 cells/well, with A1-A12 containing 0hr cells, B1-B12 containing 0hr nuclei, and C1-D12 containing 72hr nuclei (Fig. S2A), before *in situ* reverse transcription and annealing of barcode 1+linker on a thermocycler (Bio-Rad T100). After RT, cells were pooled using a multichannel pipette into a 15 mL tube, spun down at 4°C for 5 minutes at 1000 x g, and resuspended in 1 mL of Resuspension Buffer (Parse Biosciences, SB2001). Using a basin and multichannel pipette, cells were distributed in 96 wells of the Round 2 ligation barcoding plate for the *in situ* barcode 2+linker ligation. Next, cells were pooled, filtered through a 40 μm strainer, and redistributed into 96 wells of the Round 3 ligation barcoding plate for the *in situ* barcode 3+UMI+Illumina adapter ligation. After a final pooling and filtration through a 40 μm strainer, cells were counted using a hemocytometer and distributed into 7 sub-libraries: 6 sub-libraries with 9,000 cells each, and 1 sub-library with 1,000 cells. The cells in each sub-library were lysed and libraries were cleaned with AMPure XP beads (Beckman Coulter, A63881), then the single-cell barcoded cDNA underwent template switching and amplification. Importantly, we increased the number of cycles for the 1,0000-cell library to 20 cycles rather than 18 in order to increase the yield of single-cell barcoded cDNA for use in Illumina library preparation (50 ng) while having enough leftover cDNA for PacBio library preparation (500 ng). The cDNA was cleaned using AMPure XP beads and quality checked using an Agilent Bioanalyzer before proceeding to Illumina and PacBio library preparation.

### Preparation of Illumina scRNA-seq libraries using Split-seq and sequencing

All 7 sub-libraries were fragmented, size-selected using AMPure XP beads, and Illumina adapters were ligated. The cDNA fragments were cleaned again using beads and amplified, adding the fourth barcode and P5/P7 adapters, followed by a final bead-based size selection and quality check with a Bioanalyzer. Libraries with 5% PhiX spike-in were loaded at 2.1 pM and sequenced to an average depth of 51 million reads per 9,000-cell library and 70 million reads for the 1,000 cell library using an Illumina NextSeq 500 with paired-end run configuration 74/86/6/0.

### Preparation of PacBio scRNA-seq library and sequencing

The PacBio library was prepared using 500 ng of amplified, single-cell barcoded cDNA with the SMRTbell Template Prep Kit (PacBio, 100-938-900) according to the manufacturer’s protocol for sequencing on a Sequel II. The 1,000-cell library was sequenced using 2 SMRTcells (PacBio, 101-008-000) for a sequencing depth of 5,764,421 full-length non-chimeric reads.

### Preparation of bulk PacBio libraries and sequencing

We extracted RNA from two replicates of C2C12 0hr samples and 72hr samples using the RNA-easy kit (Qiagen, 74104). cDNA synthesis and library preparation using the SMRTbell Template Prep Kit (PacBio, 100-938-900) were performed as described on the ENCODE portal (https://www.encodeproject.org/documents/77db752f-abf7-4c93-a460-510464134f52). We sequenced one SMRT cell per replicate on the Sequel II platform.

### Preparation of snATAC-seq libraries using Bio-Rad technology and sequencing

The single nucleus ATAC-seq experiment was performed using the SureCell ATAC-Seq Library Prep Kit (Bio-Rad, 17004620) following the manufacturer’s protocol for the OMNI-ATAC version (45). Cells at 0hr differentiation or 72hr differentiation timepoints in one 10 cm plate per biological replicate were lifted as previously described and washed twice in cold 1X PBS + 0.1% BSA. All 0hr replicates and some 72hr replicates were filtered through a 40 μm strainer (2 technical replicates, 1 biological replicate; 2 technical replicates of 72hr samples were not filtered), then counted and assessed for viability. 300,000 cells with >90% viability per biological replicate were lysed with cold OMNI-ATAC lysis buffer on ice for 3 minutes and washed out with cold ATAC-Tween buffer, at which point non-filtered 72hr nuclei were filtered through a 40 μm strainer, then spun down at 500 RCF for 10 min at 4°C. Nuclei were resuspended, counted, and confirmed to be single-nucleus suspensions under a microscope, then 60,000 nuclei per biological replicate were tagmented at 37°C for 30 min in a ThermoMixer with 500 RPM mixing. The microfluidics-based ddSEQ Single-Cell Isolator was used to stream tagmented nuclei in an amplification reaction mix with barcoded beads to isolate single nuclei in nanodroplets with one or more barcodes. Tagmented cDNA was barcoded and amplified, then nanodroplets were broken and libraries cleaned with AMPure XP beads before a second amplification of barcoded fragments and final bead-based cleanup. A Bioanalyzer was used to verify library quality before loading at 1.5 pM and sequencing to an average depth of 122 million reads per library using an Illumina NextSeq 500 with paired-end run configuration 118/40/8/0 and custom sequencing primer.

### Validation of transcript expression with RNAscope

Myoblasts were grown to 90-100% confluency in flasks mounted on slides (Thermo Scientific, 170920) then differentiated over 3 days as previously described. The flasks were removed and slides were rinsed in 1X PBS, followed by fixation in 10% neutral buffered formalin (Sigma-Aldrich, HT501128) for 30 minutes at room temperature. Following the manufacturer’s protocol for cultured adherent cells, we rinsed the slides in 1X PBS, then incubated in 50%, 70%, and 100% ethanol for 5 minutes each (46). Slides were stored submerged in 100% ethanol at -20°C in 50 mL falcon tubes. To rehydrate, slides were incubated in 70% and 50% ethanol for 2 minutes each, then in 1X PBS for 10 minutes. A hydrophobic barrier was drawn around the edges of the slide (Vector Laboratories, H-4000), then the cells were permeabilized with 1:15 diluted protease III (ACDBio, 322340) for 10 minutes at room temperature in a humidity control tray (ACDBio, 310012). Following the manufacturer’s protocol for the RNAscope HiPlex12 kit (ACDBio, 324100/324140), probes for genes of interest were mixed and hybridized for 2 hours at 40°C in a HybEZ II hybridization oven (ACDBio, 321710/321720), then the signal was amplified over 3 rounds of 30 minute incubations at 40°C in the oven (47). We then proceeded to fluorophore hybridization and imaging over four rounds of three channels per round (GFP, RFP, and Cy5) plus DAPI (48). An EVOS FL Auto 2 with programmable stage was used to automatically image slides at 40X magnification.

### Preprocessing of LR-Split-seq data

Raw PacBio reads were processed into circular consensus reads using the ccs software from the SMRT analysis software suite (parameters: **--skip-polish --min-length=10 -- min-passes=3 --min-rq=0.9 --min-snr=2.5**) (https://github.com/PacificBiosciences/ccs). The Split-seq adapters were identified and removed using Lima (v2.0.0) (parameters: **--ccs --min-score 0 --min-end-score 0 -- min-signal-increase 0 --min-score-lead 0**) (https://github.com/pacificbiosciences/barcoding/). Reads were then processed with IsoSeq3’s Refine (v3.4.0) to yield full-length non-chimeric reads (https://github.com/PacificBiosciences/IsoSeq). As around half of our reads are primed using random hexamer priming, polyA tails were not required nor removed for this step. Reads were then demultiplexed for their Split-seq barcodes using a custom script (https://github.com/fairliereese/LR-splitpipe) by first detecting the spacer sequences between barcodes and using these as start and end points for the barcodes. Barcodes were corrected to those that were within an edit distance of 3 of the predetermined list of barcodes used for each round of barcoding. The resultant reads were then filtered on which combinations of barcodes were also seen in the Illumina single cell/nucleus RNA-seq data, which yielded 567 of the 568 cells that passed QC in the Illumina data (Fig. S2B). The reads were then trimmed of their barcodes to facilitate mapping, and cell identity barcodes were recorded. The reads were mapped using Minimap2 (v2.17-r94) (**-ax splice:hq -uf --MD**) (49) and the mm10 reference mouse genome, corrected for long-read sequencing artifacts with TranscriptClean (**--canonOnly --primaryOnly**) (50). We then used TALON (development branch on GitHub) (**--cb**) to annotate each read to its transcript or origin using the GENCODE vM21 reference (24). We filtered for reproducible novel NIC and NNC transcript models for those that were seen in 4 or more sub-cells (Fig. 1K, S1D).

### Comparing priming strategies and sample types in LR-Split-seq data

The priming strategy of each read was determined by examining the barcode for the first round of Split-seq. Reads were separated out by priming strategy and by cell. For sample comparisons, the oligo-dT and random hexamer primed reads from each cell were merged to create the final cell, then separated out by sample.

### Comparing bulk long-read to long-read Split-seq

To enable this comparison, we re-ran the bulk and single-cell data through TALON using the same database so that novel transcripts would have the same IDs across the bulk and the single-cell. For the bulk novel transcript models, filtering was done using talon_filter_trasncripts, requiring a novel transcript model to be reproducible in at least 2 of the bulk replicates with at least 5 copies. For the single-cell, filtering was done that required novel transcript models to be reproducible in at least 4 sub-cells.

### Single-cell processing of LR-Split-seq data

Oligo-dT and random hexamer primed reads from each cell were merged to create the final cells. Gene-level cells and nuclei were further filtered for those that had ≥500 reads per cell/nucleus using Scanpy (v1.4.6) (51) and for those that, in the corresponding Illumina data, had <200,000 reads, <20% mitochondrial reads, and >500 genes (done in Seurat as detailed in the Processing of short read scRNA-seq data section) (all on a per cell/nucleus basis); yielding a final total of 464 single cells and nuclei. Dimensionality reduction, construction of the UMAP, and Leiden clustering were all performed using Scanpy, yielding 7 clusters (Fig. 2D).

### Isoform switching gene testing

Testing for isoform switching in LR-Split-seq data was performed as in Joglekar et. al., 2021 (4). Tests were performed on the LR-Split-seq Leiden clusters for MB nuclei vs. *Myog^hi^* nuclei (clusters L1-L2 vs. L6-L7) and for MB nuclei vs. *Pax7^hi^* (clusters L1-L2 vs. L4-L5). Genes with significant isoform switching were required to have a corrected p-val ≤0.05 and a change in percent isoform usage per condition of ≥10, and a minimum number of reads per gene per tested condition of 10.

### Processing bulk long-read data

Bulk PacBio data was processed following the ENCODE Long Read RNA-Seq Analysis Protocol for Mouse Samples (v.1.0) for CCS, Lima, refine and TranscriptClean steps (https://www.encodeproject.org/documents/a84b4146-9e2d-4121-8c0c-1b6957a13fbf). A TALON database was initialized using mm10 GENCODE v21 GTF with SIRV set 3 and ERCCs included. Reads output from TranscriptClean were labeled with the corresponding fasta reference. TALON was run **(--cov 0.9 --identity 0.8)**. Filtering novel transcript models was done using TALON’s talon_filter_transcripts module, requiring a novel transcript model to be reproducible across biological replicates, and appear 5 times in each replicate, as well as display a lack of internal priming evidence (**-- minCount 5 --minDatasets 2 --maxFracA 0.5**). Transcript abundances were determined using talon_abundance.

### Processing of short read Split-seq data

After initial demultiplexing of the 7 sublibraries (6x 9,000-cell sublibraries and 1 1,000-cell sublibrary), Parse Bioscience’s split-pipe v0.7.6 software was used to deconvolute reads into single cells, map to mm10 using STAR (v. 2.6.0c), annotate using GENCODE vM21, and filter using a UMI cutoff determined by knee plots (Fig.S2B, S2D) (52). The remaining cells were further filtered in Seurat (v. 3.2.2) by <20% mitochondrial reads, < 200,000 counts, and >500 genes per cell/nucleus (Fig. S2C, S2E) (53). The resulting 464 cells with both short and long reads and 36,405 cells with short-read data only were analyzed using Velocyto (v.0.1.17) (25). 55% of counts from 0hr cells, 46% of counts from 0hr nuclei, and 37% of counts from 72hr nuclei were spliced out of the total number of spliced and unspliced counts. After loading the loom file back into Seurat with the ReadVelocity function from the SeuratWrappers package, SCTransform (v. 0.3.1) was used to regress percent mitochondrial reads, number of genes, and sublibrary, followed by UMAP dimensionality reduction (54, 55). Clustering using the Leiden algorithm (v. 0.8.0) resulted in 20 clusters (56). Differentially expressed genes per cluster were found using Seurat’s FindAllMarkers function (**only.pos = TRUE, min.pct = 0.1, logfc.threshold = 0.1**) then further filtered by FDR < 0.01.

### Processing of snATAC-seq data

After demultiplexing the 8 snATAC-seq libraries (3 0hr, 5 72hr samples), Bio-Rad’s dockerized ATAC-seq analysis toolkit (v.1.0.0) was used to recover barcodes/UMIs, align reads with BWA, filter and deconvolute barcodes, perform quality control by UMI thresholding, and call peaks with MACS2 (Fig. S3A) (57, 58, 59). A custom script (https://github.com/fairliereese/lab_pipelines/tree/master/sc_atac_pipeline) that takes in the combined peaks file, QC-passing barcode list, and mapped reads was used to generate peaks-by-cells counts matrices as csv files for each library. In addition, the annotated bam files were converted to fragment files using scATAC-pro’s simply_bam2frags.pl script, which are bed-like matrices containing chromosome, start, stop, cell ID, and number of fragments contained in the region (60). Further QC cutoffs consisted of a TSS enrichment score > 6, > 5,000 counts, and < 20,000 counts per nucleus (Fig. S4B). TSS enrichment is calculated in Signac (v. 1.0.9004) following the definition by ENCODE (https://www.encodeproject.org/data-standards/terms/). Signac was used to normalize binarized peaks-by-cells counts matrices by term frequency inverse document frequency (TF-IDF) followed by singular value decomposition and UMAP dimensionality reduction (37). The Leiden algorithm (v. 0.8.0) was used to resolve 18 clusters (Fig. 4A). UCSC Genome Browser tracks were generated by splitting the snATAC bam file by cluster using the sinto package (https://github.com/timoast/sinto) and creating bigwig tracks using deeptools (61). Differentially accessible peaks per cluster were found using Seurat’s FindAllMarkers function (**only.pos = TRUE, min.pct = 0.1, logfc.threshold = 0.5**) then further filtered by FDR < 0.05. The marker peaks were grouped by MB (A1-A7), *Pax7*^hi^ (A8-A15), and *Myog^hi^* (A16-A17) and processed using GREAT with mm10 whole genome background and associating peaks with the single nearest gene within 50kb (62). P values for the binomial test are reported in the text.

### Integration of short-read Split-seq and snATAC-seq data

Signac’s FindTransferAnchors function was implemented with all 36,869 Split-seq cells as the reference set and all 23,525 snATAC-seq nuclei as the query set, with canonical correlation analysis (CCA) used as the dimensional reduction method (38). The TransferData function was used to carry over Split-seq labels “0hr” or “72hr” in one analysis (Fig. S4D, left panel) and labels “MB”, “*Pax7*^hi^”, and “*Myog*^hi^” in another analysis (Fig. S4D, right panel).

### Identification of TSSs from long-read data

For both LR-Split-seq and bulk separately, bam reads were filtered for those that were annotated by TALON as belonging to the known, novel in catalogue (NIC), novel not in catalogue (NNC), and prefix-ISM novelty categories as the starts of reads belonging to these novelty categories are more likely to come from a true 5’ end. TSSs were called on the filtered bams using the ENCODE PacBio TSS caller (https://github.com/ENCODE-AWG/tss-annotation/blob/master/long_read/pacbio_to_tss.py) (**--window-size=50 --raw-counts - -expression-threshold=0)**, yielding a bed entry for each TSS consisting of a wide peak, narrow peak, and a summit for each TSS. Resultant Split-seq TSSs were filtered first by requiring each one to be supported by at least 2 reads, and subsequently on the gene level, where each called TSS was required to have a number of reads >10% of the number of reads that supported the most highly expressed TSS for the same gene. Bulk TSSs were similarly filtered except using a threshold of >5%.

### Identification of TESs from long-read data

Similarly, for both LR-Split-seq and bulk data separately, bam reads were filtered for those annotated as belonging to the known, novel in catalogue (NIC), novel not in catalogue (NNC), and suffix-ISM novelty categories as the ends of reads belonging to these novelty categories are more likely to come from a true 3’ end. TESs were called on the filtered bam using the same ENCODE PacBio TSS caller (https://github.com/ENCODE-AWG/tss-annotation/blob/master/long_read/pacbio_to_tss.py) (**--window-size=50 --raw-counts - -expression-threshold=0 --tes)**, yielding a bed file the same format as the TSS file. The TESs were then filtered by requiring each to be supported by at least 2 reads and have a number of reads >80% of the number of reads that supported the most highly-expressed TES for the same gene.

### Processing C2C12 CAGE data

CAGE data was downloaded from GEO accession GSE21580 (40). Wig files corresponding to CAGE data from days 0 and 9 of C2C12 differentiation were converted to bed format using bedops wig2bed (63) and lifted over from the mm9 genome to the mm10 genome using UCSC’s liftOver tool **(-minMatch=0.95)** (64). Resultant bed peaks were concatenated.

### Processing C2C12 PolyA-seq data

PolyA-seq data was downloaded from GEO accession GSE62001 (65). Entries in the provided expression matrix were filtered for those belonging to the “C2C12.Pro” (proliferating C2C12) and “C2C12.Diff” (4-day differentiation C2C12) categories. The data was then converted into bed format using a custom script and lifted over from mm9 to mm10 using the UCSC liftOver tool **(-minMatch=0.95)** (64).

### Intersecting TSSs with validation datasets

A combined TSS validation bed file was made using the proximal enhancer and promoter ENCODE cCREs (41), GENCODE vM21 TSSs (66), our snATAC-seq pseudobulk peaks, and CAGE peaks (40). The filtered TSSs for both bulk (22,938) and LR-Split-seq (23,996) were intersected with the combination bed file using bedtools intersect with default parameters, meaning minimum of 1bp overlap between the TSSs and the combined validation set (2,057,291 regions) (67).

### Intersecting TESs with validation datasets

A combined TES validation bed file was made using our snATAC-seq pseudobulk peaks and polyA-seq peaks (65). Similar to TSS validation, bedtools intersect with default overlap settings (1bp) was used to determine the number of overlaps between our filtered TESs for both bulk (14,120) and LR-Split-seq (12,521) and the combined validation set (205,853 regions).

### snATAC-seq and TSS integration

TSS regions identified in LR-Split-seq in bed format were used to calculate activity at each TSS through the Signac interface. Normalized expression values and normalized TSS activity values were averaged across the three groups of cells (MB, *Pax7^hi^*, and *Myog^hi^*) and a pseudocount of 1 was added to each TSS. Fold change in expression and activity separately was calculated by dividing the TSS values of one group by another group, such as *Myog*^hi^/MB. The log2 fold change for each TSS was then plotted for both expression (x-axis) and activity (y-axis), revealing TSSs with chromatin profiles and expression in agreement at the upper right and bottom left sectors. Twice the standard deviation of each dataset is indicated by black dashed lines. (Fig. 4F-4H).

### LR-Split-seq TSS quantification and differential TSS testing

TSS expression was quantified from the LR-Split-seq data starting from the TALON read annotation file (Table S1), which tracks the start and end coordinates of every read. Read starts were converted to a read start bed file and expanded to include ±25 bp from the true start. Finally, the read start bed was intersected using bedtools with the filtered LR-Split-seq TSSs (Table S8), requiring at least 1 bp of overlap. The number of reads per TSS was then computed by counting all of the reads assigned to each TSS. Testing on the TSS level for the LR-Split-seq data was performed as in Joglekar et. al., 2021 (4). Tests were performed on the LR-Split-seq Leiden clusters for MB nuclei vs. *Myog^hi^* nuclei (clusters L1-L2 vs. L6-L7) and for MB nuclei vs. *Pax7^hi^* (clusters L1-L2 vs. L4-L5). Genes with significant TSS switching were required to have a corrected p-val ≤0.05 and a change in percent isoform usage per condition of ≥10, and a minimum number of reads per gene per tested condition of 10.

## Supporting information

Supupplemental Tables

## Acknowledgements

We would like to thank Melanie Oakes at UC Irvine Genomics High-Throughput Facility (GHTF) for her help with PacBio sequencing. This work was supported in part by grants from the National Institutes of Health (UM1HG009443) to A.M. and B.J.W.

## Notes

### Competing Interest Statement

The authors have declared no competing interest.

