## Supplementary Figures for "Mapping and modeling the genomic basis of differential RNA isoform expression at single-cell resolution with LR-Split-seq"

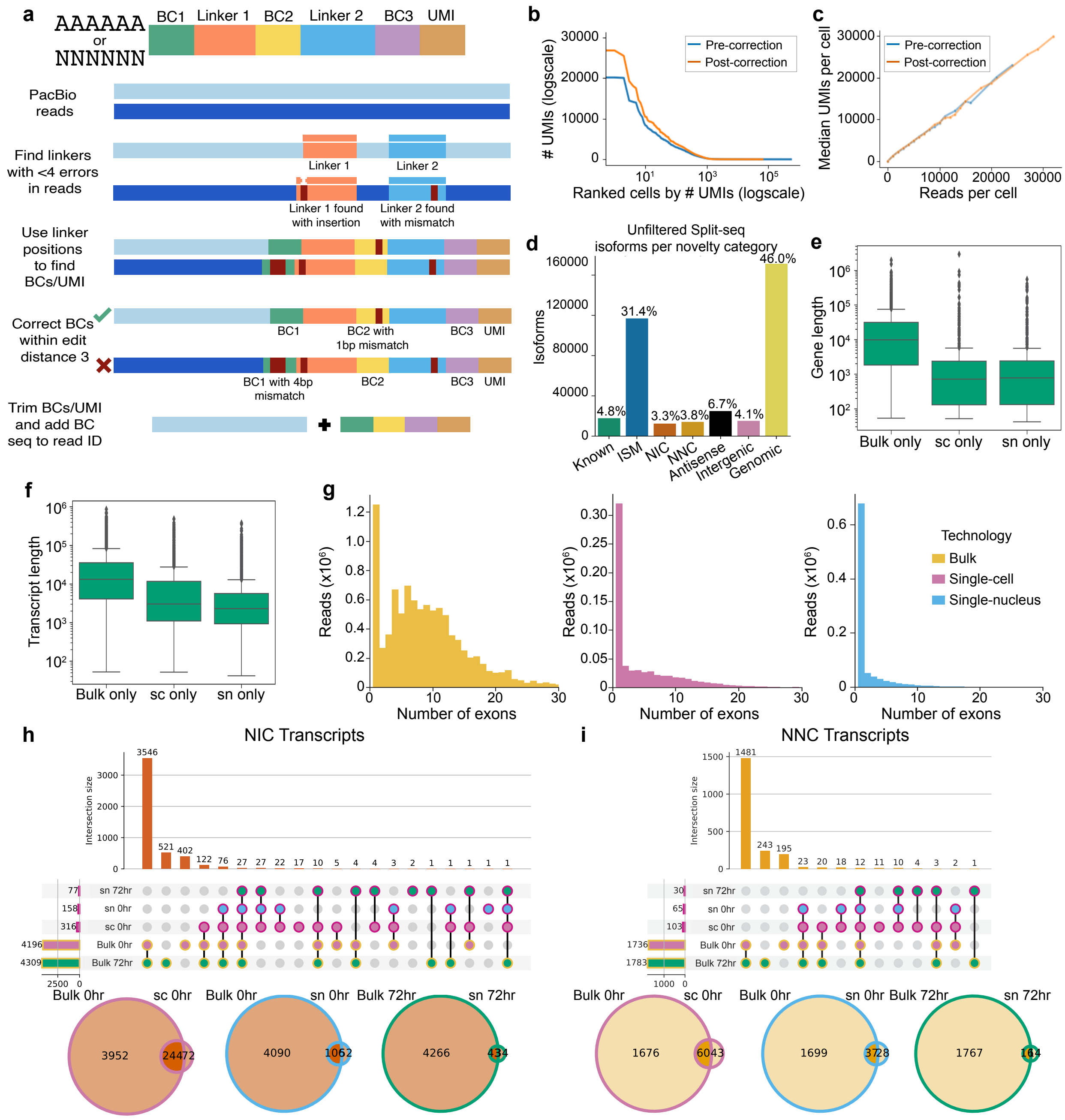

**Supplemental Fig. 1: LR-Split-seq preprocessing, QC, and additional analysis. a**, Schematic diagram of LR-Split-seq demultiplexing strategy. **b**, UMI per ranked barcode plots before and after barcode correction (both axes log scaled). **c**, Median number of UMIs per cell binned by reads per cell before and after barcode correction. **d**, Unfiltered isoforms per novelty category across all cells in LR-Split-seq data. **e**, Gene lengths of annotated genes detected in bulk only, single-cell only, and single-nucleus only (log scale). **f**, Transcript lengths of annotated transcripts detected in bulk only, single-cell only, and single-nucleus only (log scale). **g**, Distribution of number of exons in bulk long reads (yellow), single-cell long reads (pink), and single-nucleus long reads (blue). **h**, Upset plot of novel in catalog (NIC) transcripts that passed filtering found in bulk data compared to single cell data across all samples. Bars on the left indicate set size, circles indicate various combinations of samples, and bars on top indicate the number of genes found in each combination. Outline colors indicate technology (bulk in yellow, single-cell in magenta) and fill colors indicate sample type (72hr nuclei in green, 0hr nuclei in blue, and 0hr cells in pink for single-cell data; 72hr in green, 0hr in pink for bulk data). Box plots above indicate gene length distribution for each intersection. Venn diagrams below summarize the overlaps between bulk (left) and single-cell or single-nucleus (right), for each sample type. Sample type is indicated by outline color. **i**, Upset plot and Venn diagrams of novel not in catalog (NNC) transcripts that passed filtering found in bulk data and single-cell data.

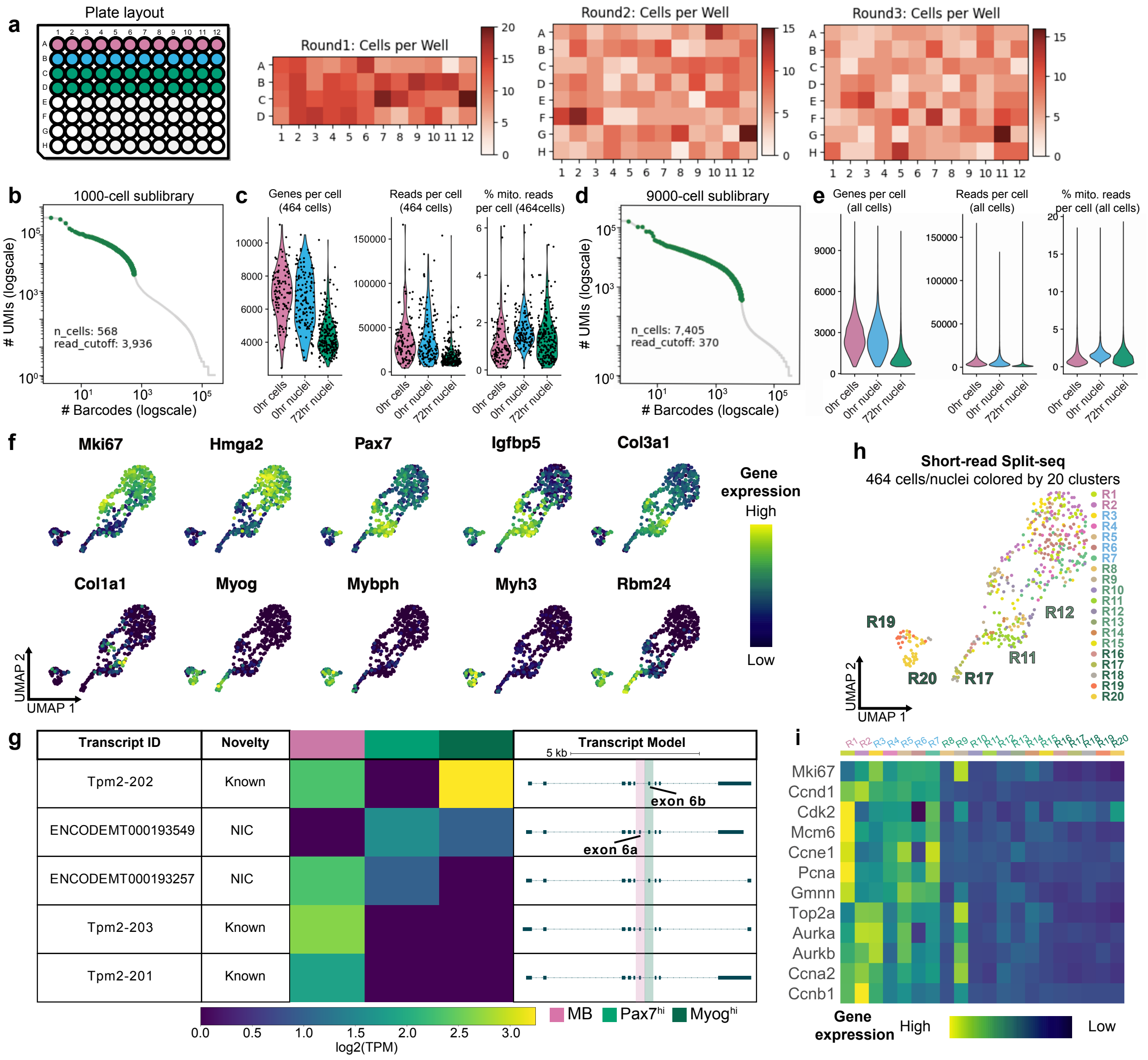

**Supplemental Fig. 2: Short- and long- read Split-seq QC and additional analysis.** **a**, Schematic of sample type per well in the first round of barcoding (pink = 0hr cells, blue = 0hr nuclei, and green = 72hr nuclei). Panels to the right show the number of cells per well across each round of barcoding for a 9,000-cell sublibrary. **b**, UMI per cell knee plots for the 1,000-cell sublibrary sequenced with both long and short reads indicating a threshold of 3,936 reads per cell, leaving 568 cells before additional QC. **c**, Violin plots of scRNA-seq QC metrics after filtering for the 464 cells only. **d**, An example knee plot for a 9,000-cell sublibrary indicating a threshold of 370 reads per cell, leaving 7,405 cells before additional QC. **e**, Violin plots of scRNA-seq QC metrics after filtering for all cells. **f**, Distribution of marker genes within the 464-cell UMAP (dark blue = lowly expressed, yellow = highly expressed). **g**, Gene report made by Swan for Tpm2. Relative expression of each isoform, separated by 0hr MB cells, 72hr *Pax7*$^{hi}$ nuclei, and 72hr *Myog*$^{hi}$ nuclei plotted alongside the isoform's name, transcript novelty, and structure. Exons 6a and 6b, known to be alternatively spliced during C2C12 differentiation, are highlighted. **h**, UMAP of 464 cells with both short and long reads colored by 20 clusters derived using 36,869 short-read cells. **h**, Heatmap of cell cycle marker genes in the 20 clusters.

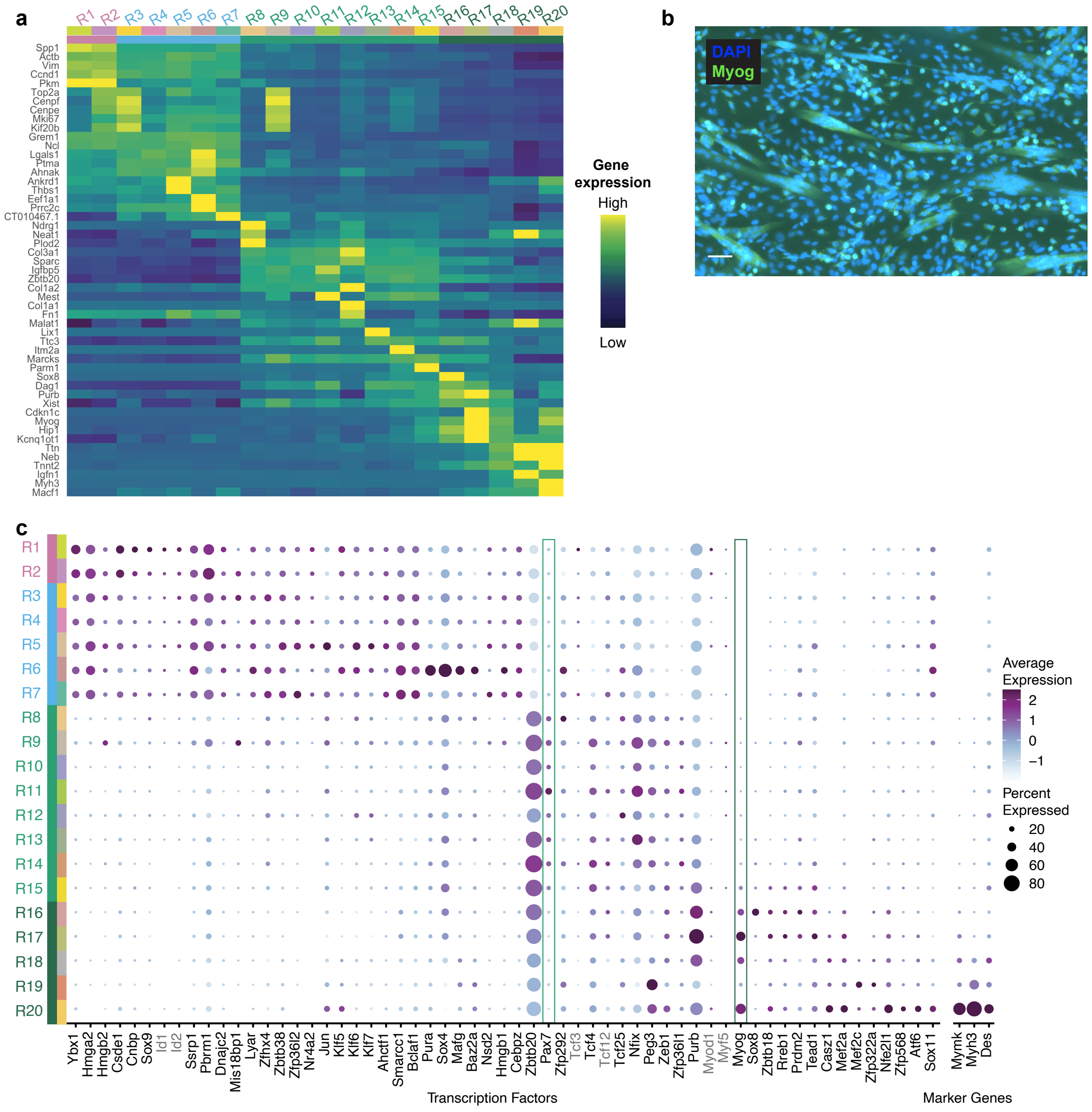


**Supplemental Fig. 3: Short-read Split-seq additional analysis. a**, Heatmap of marker genes in the 20 clusters (dark blue = low expression, yellow = high expression). **b**, Visualization of *Myog* in mononucleated cells and myotubes at the 72hr differentiation timepoint. Blue = DAPI, green = *Myog*. Scale bar: 50 μm. **c**, Dot plot of transcription factors and marker genes involved in myogenesis found from differential expression testing and/or literature. Genes that did not pass the differential expression threshold yet are of interest in the system and significantly expressed in prior classic bulk data are colored grey (*Id1*, *Id2*, *Myod1*, *Myf5*, *Tcf3,* and *Tcf12*).

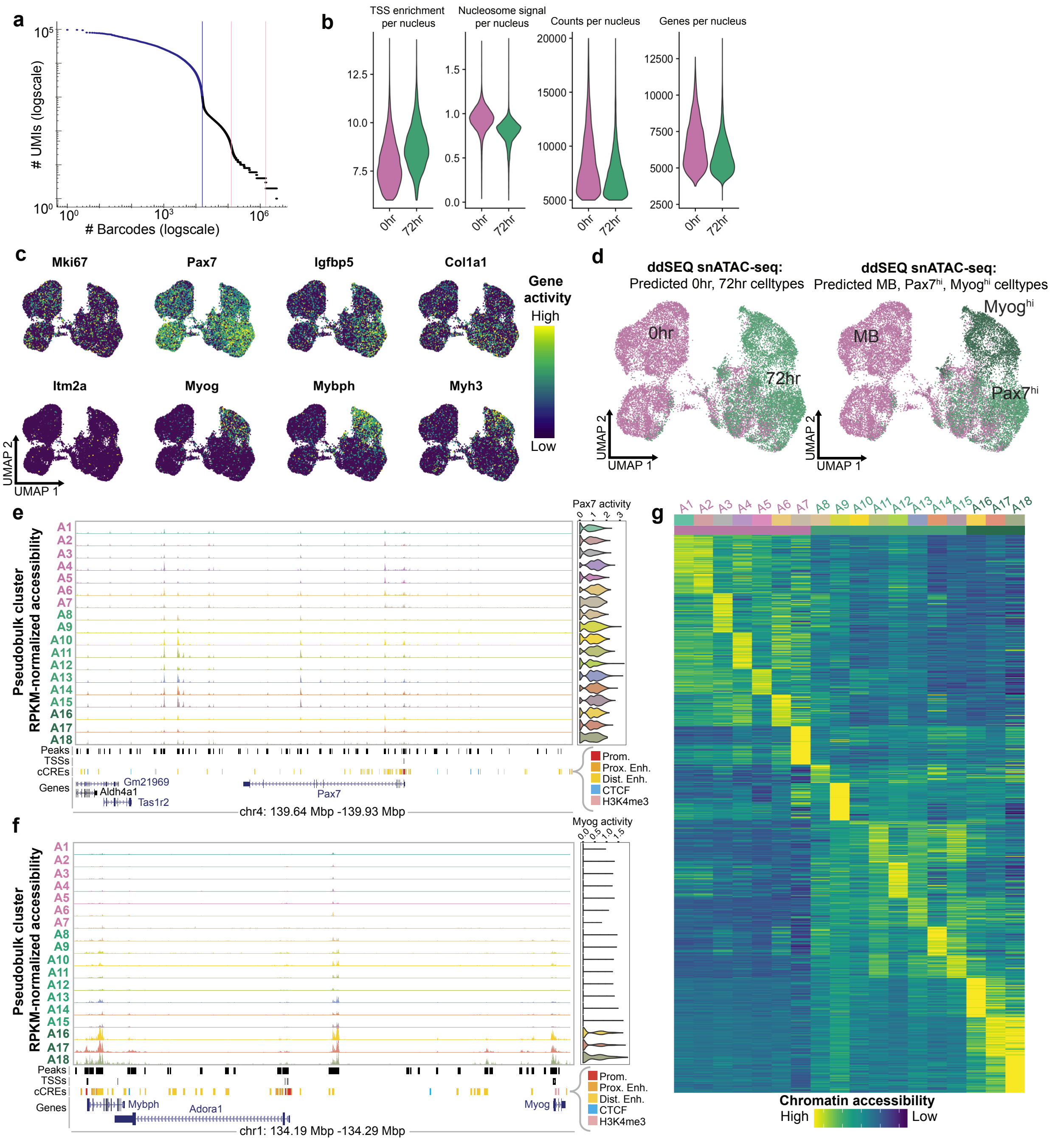


**Supplemental Fig. 4: Additional analysis/QC of snATAC-seq.** **a**, UMI per barcode knee plot for an example snATAC-seq library (0hr, 6,782 nuclei). **b**, Violin plots of snATAC-seq QC metrics after filtering > 6 TSS enrichment, < 20,000 reads, and > 5,000 reads per nucleus. **c**, Distribution of marker genes within the UMAP colored by gene activity score (dark blue = low activity, yellow = high activity). **d**, Integration of scRNA-seq and snATAC-seq data, labeled by cell type (0hr in pink and 72hr in green on left; MB in pink, *Myog*$^{hi}$ in dark green, and *Pax7*$^{hi}$ in light green on right). **e**, Pseudobulk peaks per cluster spanning the *Pax7* locus. TSS track indicates TSSs called from LR-Split-seq data. **f**, Pseudobulk peaks spanning the *Myog* and *Mybph* loci. **g**, Heatmap of top 50 marker regions in the 18 snATAC-seq clusters (dark blue = low accessibility, yellow = high accessibility).

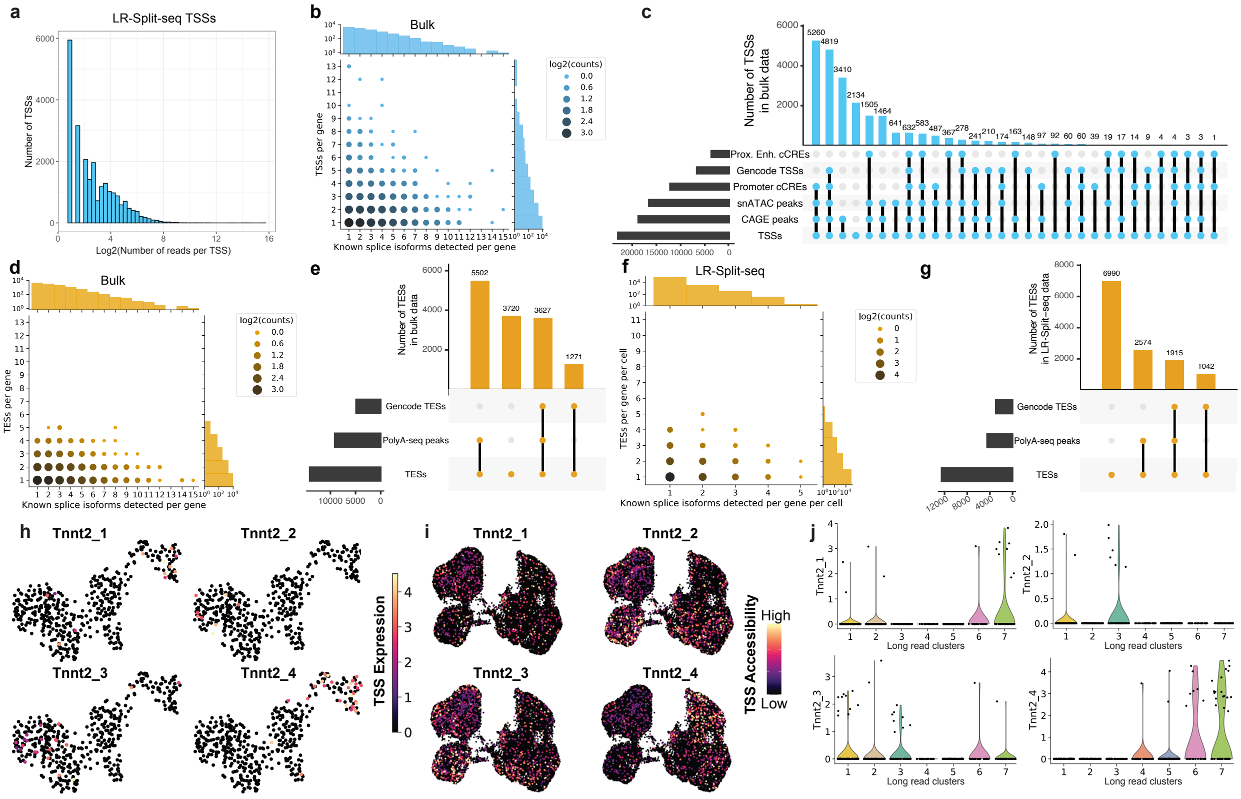


**Supplemental Fig. 5: Identification and validation of TSSs/TESs from long-read data. a**, Histogram of number of LR-Split-seq reads supporting each TSS. **b**, Bubble plot of the number of distinct exon combinations (splice isoforms) detected per gene compared to the number of distinct TSSs detected per gene in bulk data. **c**, Validation of TSSs found in bulk long-read data using 4 external datasets (ENCODE proximal enhancer and promoter cCREs, GENCODE TSSs, and CAGE peaks) and our snATAC-seq pseudobulk peaks. **d**, Bubble plot of the number of distinct exon combinations (splice isoforms) detected per gene compared to the number of distinct TESs detected per gene found in long-read bulk data. **e**, Validation of TESs found in bulk long-reads using GENCODE TESs and polyA-seq data. **f**, Bubble plot of splice isoforms per gene per cell compared to TESs detected per gene per cell found in LR-Split-seq. **g**, Validation of TESs found in LR-Split-seq. **h**, LR-Split-seq TSS expression for the 4 identified *Tnnt2* TSSs. **i**, snATAC accessibility for the 4 identified *Tnnt2* TSSs. **j**, Violin plots of TSS expression per long read cluster for the 4 identified *Tnnt2* TSSs.
